# Zebrafish studies on the vaccine candidate to COVID-19, the Spike protein: Production of antibody and adverse reaction

**DOI:** 10.1101/2020.10.20.346262

**Authors:** Bianca H Ventura Fernandes, Natália Martins Feitosa, Ana Paula Barbosa, Camila Gasque Bomfim, Anali M. B. Garnique, Francisco I. F. Gomes, Rafael T. Nakajima, Marco A. A. Belo, Silas Fernandes Eto, Dayanne Carla Fernandes, Guilherme Malafaia, Wilson G. Manrique, Gabriel Conde, Roberta R. C. Rosales, Iris Todeschini, Ilo Rivero, Edgar Llontop, German G. Sgro, Gabriel Umaji Oka, Natalia F Bueno, Fausto K. Ferraris, Mariana T. Q. de Magalhaes, Renata J. Medeiros, Juliana M. M Gomes, Mara de Souza Junqueira, Katia Conceição, Letícia G. Pontes, Antonio Condino-Neto, Andrea C. Perez, Leonardo J. G. Barcellos, Jose Dias Correa junior, Erick G. Dorlass, Niels O. S Camara, Edison Luiz Durigon, Fernando Q. Cunha, Rafael H. Nóbrega, Glaucia M. Machado-Santelli, Chuck Farah, Flávio P Veras, Jorge Galindo-Villegas, Leticia Costa-Lotufo, Thiago M. Cunha, Roger Chammas, Cristiane R. Guzzo, Luciani R Carvalho, Ives Charlie-Silva

## Abstract

Establishing new experimental animal models to assess the safety and immune response to the antigen used in the development of COVID-19 vaccine is an imperative issue. Based on the advantages of using zebrafish as a model in research, herein we suggest doing this to test the safety of the putative vaccine candidates and to study immune response against the virus. We produced a recombinant N-terminal fraction of the Spike SARS-CoV-2 protein and injected it into adult female zebrafish. The specimens generated humoral immunity and passed the antibodies to the eggs. However, they presented adverse reactions and inflammatory responses similar to severe cases of human COVID-19. The analysis of the structure and function of zebrafish and human Angiotensin-converting enzyme 2, the main human receptor for virus infection, presented remarkable sequence similarities. Moreover, bioinformatic analysis predicted protein-protein interaction of the Spike SARS-CoV-2 fragment and the Toll-like receptor pathway. It might help in the choice of future therapeutic pharmaceutical drugs to be studied. Based on the *in vivo* and *in silico* results presented here, we propose the zebrafish as a model for translational research into the safety of the vaccine and the immune response of the vertebrate organism to the SARS-CoV-2 virus.

## Introduction

The World Health Organization (WHO) registered, on January 30th, 2020, that the outbreak of the disease caused by the severe acute respiratory syndrome coronavirus 2 (SARS-CoV-2) constituted a Public Health Emergency of International Importance (the highest level of alert from the Organization)1. Since then, the number of Coronavirus Disease 2019 (COVID-19) cases outside China has increased significantly worldwide, resulting in the deaths of approximately 1,108,000 of the nearly 40 million infected people through October1. The current coronavirus pandemic has had drastic consequences for the world’s population, not only in terms of the public health system but also in causing a major global economic crisis. Diagnostic tests, efficient and safe vaccines, and new effective antivirals are urgently required2.

SARS-CoV-2 Spike protein is found on the surface of the virus, giving it a “crown” appearance, and binds human (Homo sapiens) Angiotensin-converting enzyme 2 (ACE2) to infect human cells7. Moreover, the Spike protein is one of the likely targetsfor vaccine production, and the antibodies against it could be used for SARS-CoV-2 antigen rapid test production. To investigate the production of specific antibodies against the Spike protein of SARS-CoV-2 in a zebrafish (Danio rerio) model, we inoculated an N-terminal region of SARS-CoV-2 Spike recombinant protein (residues 16-165) in adult female specimens. In humans, protection conferred by natural infection or passive immunization are unclear^1^. However, in teleost fish, including the zebrafish, antibodies constitute a major proportion of the functional passive immunity that is aquired maternally. Although maternal Abs are transferred to the fetus through the placenta in mammals, in almost all teleost fish, Abs are transferred to the yolk^2^. This suggests that by injecting the recombinant spike protein abundant antibodies could be obtained simply by extracting the antibodies from the eggs produced by a single adult female zebrafish. The second goal of this work was to demonstrate that the zebrafish could be a new alternative model to test preclinical vaccine candidates for COVID-19, prospecting a strategy to assess safety and toxicity for vaccine candidates. The comparison between zebrafish and human genomes revealed remarkable sequence and functional conservation of 70% genetic similarity to humans^3,4^. Zebrafish have been used as a model to study the safety of vaccines^5^ and to assess toxicology that could be correlated to human health^6,7^. Recently, the WHO (2020) prepared a document on all vaccine candidates for COVID-19 clinical trials, reporting 35 candidate vaccines in the clinical evaluation and at least 166 vaccine candidates in preclinical and clinical development. Normally, the development of a vaccine takes 10 to 15 years for conclusion. However, the case of COVID-19 meets a new pandemic paradigm and the development of the vaccine has been proposed to be reduced to 1-2 years^8^.

It is worth mentioning that before vaccine clinical tests begin, several safety protocols must be submitted with *in vitro* and *in vivo* experiments on animal models. There is a lack of information regarding the immune response of the organism to SARS-CoV-2, including animal models to study it^9^. Although zebrafish do not have lungs as humans do, the present study shows similar inflammatory responses observed in severe cases of COVID-19 patients that could be considered when investigating human responses to the virus.

In the global task to develop the vaccine and possible therapeutic approaches for COVID-19, several animal models have been proposed, such as mice^10^, hACE2 transgenic mice^11^, alpaca^12^, golden Syrian hamsters, ferrets, dogs, pigs, chickens, and cats^9^, and species of non-human primates^10^. Recently, three reports have described the production of equine neutralizing antibodies against SARS-CoV-2^13,14^. A study by Deng and collaborators analyzed serum samples from 35 animal species for the detection of specific antibodies against SARS-CoV-2^15^. Despite this wide search for candidate animal models, so far only two references promote the zebrafish model on this regard confirming the innovative and pioneer characteristics of our study^16,17^.

Here, female zebrafish individuals injected with a N-terminal fraction of SARS-CoV-2 Spike recombinant protein (residues 16-165) produced specific antibodies, and presented suggestive adverse reactions and inflammatory responses resembling the severe cases of COVID-19 human patients. Therefore, with this work we put forward the advantage of using zebrafish as a model for translational research on the vaccine safety and the screening of immune response against the SARS-CoV-2 virus.

## Results

### Humoral immune response of zebrafish immunized with N-terminal fraction of rSpike protein

To induce and analyze the humoral immune response, 3 peptides of full length SARS-CoV-2 Spike were generated after a pattern memorizing phagolysosomal proteolysis using the virtual proteolytic cleavage tool (Figure 1a and 1a.1). One of them, the peptide named Pep1 (Pep1, residues 16 to 22; Pep2 and Pep3 are shown in Supplemental Figure 1) (Figure 1a-a2), has been chosen because of its promising antigenic potential. It has presented a binding free energy site in protein-ligand interactions for *D. rerio* MHC II, MHC I, TCR alpha (Figure 1b). Similar results were observed when the same analysis was performed using human orthologous receptors (Figure 1b). Further analysis were carried out based on Dock analysis between Pep15 and the structure of MHC II (PDBID), MHC I (PDBID), TCR alpha (PDBID), and TCR beta (PDBID) that showed the similarity of the ligand/Pep1 interaction to the receptor-binding site (Figure 1c). After the *in silico* examination, specific pathogen free wildtype (AB SPF) adult female zebrafish were injected with a N-terminal fragment of SARS-CoV-2 Spike protein (residues 16 to 165) expressed in *Escherichia coli* with a N-terminal fusion of six histidine tag and purified from inclusion bodies, herein name rSpike, to determine whether they could produce IgM-class antibodies. In 7 days of immunization, a band corresponding to IgM in the plasma was detected using SDS-PAGE and was also and analyzed by MALDI-ToF. It was two-fold higher than the controls (Figure 1e, g). After 7 days a new immunization using rSpike was done and the IgM level remained higher than the control after 14 days being more evident in IgM of eggs (Figure 1e, g).

**Figure 1.**
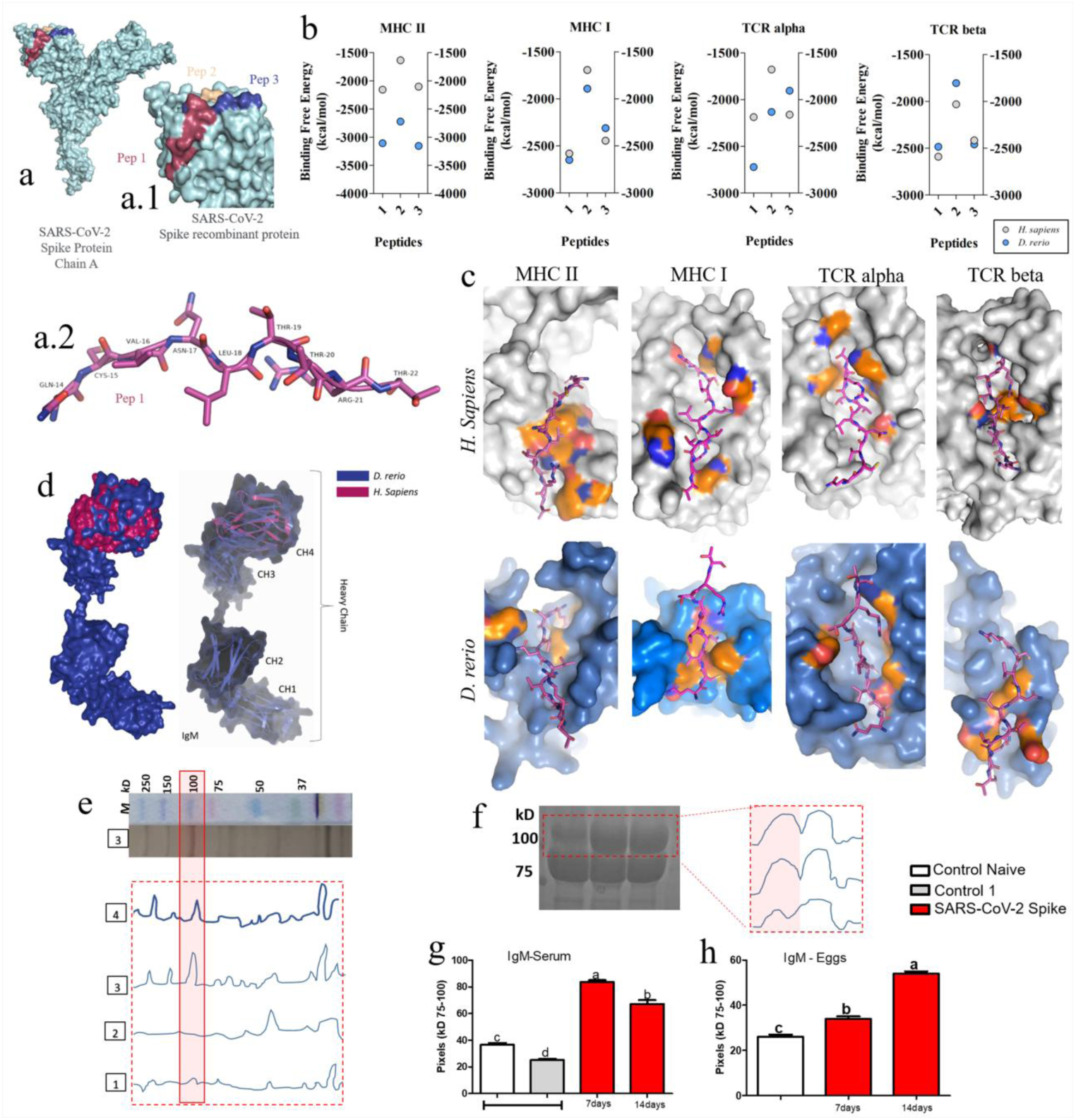
rSpike protein and its effects on the humoral immune response *in silico* and *in vivo* in zebrafish. (a-a.1) Cryo-EM structure of the SARS-CoV-2 Spike Protein (PDBID 6cs2.1, chain A) highlighting the residues 16-165 in blue (Pep1), in red (Pep2), and yellow (Pep53). (a.2) Representation of the peptide 1 (residues 14-22), oxygen, nitrogen and carbon are colored in red, blue and pink, respectively. (b) Free binding energy of SARS-CoV-2 Spike Pep1, Pep 2, and Pep 3 in complex with MHC II, MHC I, TCR alpha, and TCR beta of human (grey dots) and zebrafish (blue dots) based on docking analysis and the axis (X) represents the score of 10 (ten) possibilities of interaction between molecule-ligand and the axis (Y) compares the free binding energy it represents per kilocalorie per mol (Kcal/mol). (c) Comparison of topological location and insertion of Pep 1 in the receptor protein binding site from zebrafish (botton panel) and human (top panel) MHC II, MHC I, TCR alpha, and TCR beta. The amino acid receptor residues are shown on the protein surface in orange colors; red and blue represented by the chemical elements. (d) Structural alignment of the IgM constant/heavy chain between zebrafish and human. (e) Densitometry of 100 kDa bands from adult female serum separated by a SDS-PAGE (red colored box): M: molecular weight marker (company) and the red dotted box correspond to intensities of the bands from the SDS-PAGE of naïve female serum (box 1), IgM production from immunized zebrafish with buffer (box 2), and rSpike protein (residues 16-165) after 7 (box 3) and 14 (box 4) days. (f) Densitometry of 100 kDa bands of a SDS-PAGE gel loaded with eggs extract from naïve (1) and female injected with rSpike (residues 16-165) after 7 (2) and 14 days (3). (g, h) Graphs representation of densitometry quantification of serum (g) and egg (h) IgM levels showed in panel e and panel f, respectively, demonstrating an increase of IgM production by immunized females (red bars). Control naïve are fishes not treated, Control 1 are fishes treated with buffer and rSpike protein (spike residues 16-165).

Docking analysis showed that the zebrafish IgM chain 4 (CH4) share 43.3% sequence similarity to human IgM CH4 (Figure 1d) and might have similar potential to recognize S protein as the human antibody. In parallel, it was tested whether passive antibody transfer to the eggs occurred through immunized females. The bands corresponding to the size of IgM in unfertilized zebrafish eggs were detected in the gel (Figure 1f) and confirmed by protein analysis with MALDI-ToF. It was possible to observe an increase in IgM in eggs compared to the control after 7 days, and it was almost two-fold higher after 14 days of maternal immunization (Figure 1h).

### rSpike protein immunization of zebrafish had an impact on the survival rate

Two bioassays were carried out to analyze the toxicity of the rSpike. Although the immunized fish produced antibodies, the first injection of the rSpike generated high toxicity to the fish (Figure 2). Therefore, the assay was repeated by adding different control groups to confirm that the toxicity findings were specific to the rSpike (Figure 2). In the first bioassay, after fish immunization with rSpike, the survival rate was 78.6% during the first seven days (Figure 2). It was significant when compared to naive control and fish injected with protein buffer (control 1), where the survival rate was 100% and 90%, respectively (Figure 2). Nonetheless, after a second immunization, the rSpike immunized group maintained the plateau survival rate, with no statistical significance between the groups for the relative risk of death (Figure 2).

**Figure 2.**
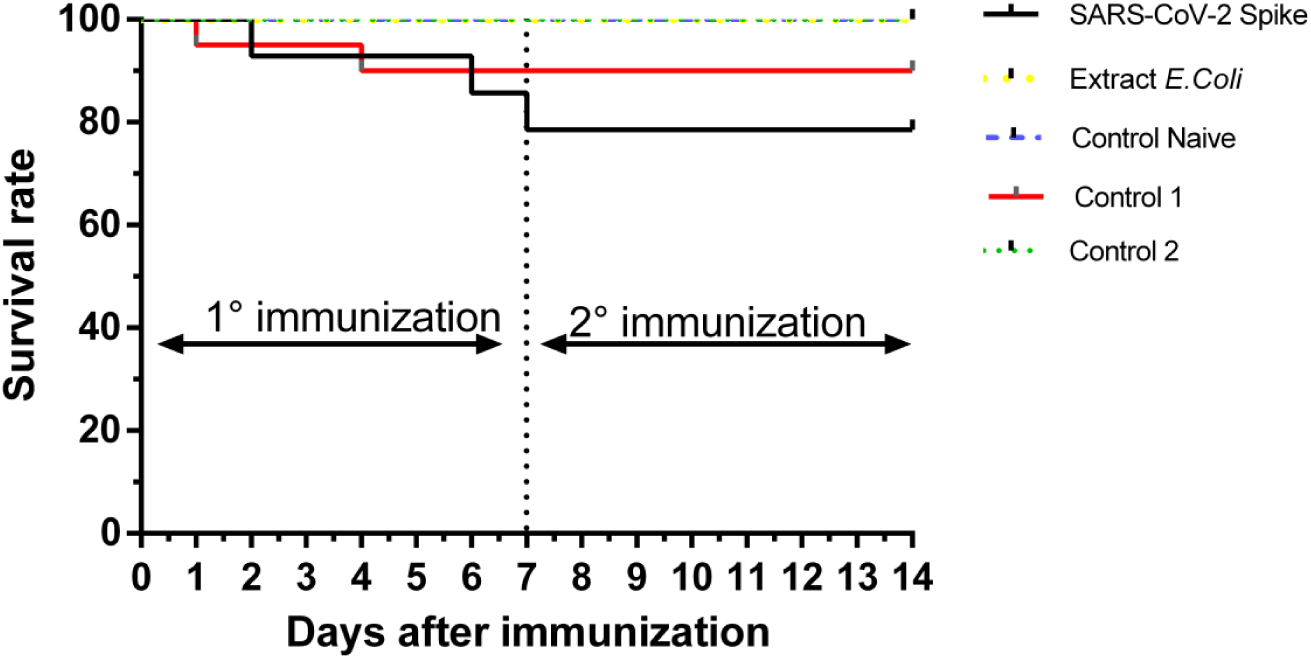
rSpike protein injection is toxic to adult female zebrafish. Graph of survival rate and days after immunization. Kaplan-Meier cumulative probability curve indicating survival rate of zebrafish after two immunization with different protein samples. Females were injected either with rSpike protein, extract of lysed *E*.*coli* cells, buffer presented the rSpike protein (control 1), naïve control (not immunized), or a mix of two recombinant protein: PilZ protein from *Xanthomonas citri* and N-terminal part of LIC_11128 from *Leptospira interrogans* Copenhageni (control 2). Each group was performed using adult female fishes.

Therefore, a second assay was conducted by adding different control groups in order to confirm that the toxicity findings were specific to the rSpike, and related to the presence of any antigen. The Kaplan-Meier survival analysis confirmed rSpike injection presented a lower survival rate compared to the two previous controls used (Control naïve and protein buffer) and also compared to females injected with *E. coli* extract or a culture medium mixed of two purified recombinant proteins (PilZ protein from *Xanthomonas citri*, and a N-terminal fragment of LIC_11128 from *Leptospira interrogans* Copenhageni) (Control 2) (Figure 2). The survival rate was maintained after the second immunization for the next seven days. The relative risk of death in the period studied between the groups was significant (chi square = 79.70; p <0.0001).

### rSpike protein produced an inflammatory response and critical damage in different tissues of adult zebrafish

In order to verify the occurrence of sublethal effects of the rSpike on treated zebrafish, histopathological analysis of different organs, including brain, gonads, heart, kidney, liver, spleen, among others, was performed in female fishes used in the immunization protocol described in material and methods. Animals that died during the immunization experiment were excluded from the analysis. In general, it was observed several morphological alterations compatible with an undergoing inflammatory process in many tissues. Markedly, brain obtained from treated fishes showed an intense inflammatory infiltrate with presence of many macrophages after 7 days (Figure 3c) and an intense mononuclear infiltrate after 14 days (Figure 3d,e). Histopathological analysis of the female reproductive tissue showed ovarian stroma with abundant and disorganized extracellular matrix (Figure 3g). Follicular development showed alterations such as atresia among oocytes at primary growth and cortical alveolus stages (Figure 3g). Moreover, dense inflammatory infiltrates are commonly seen in the ovarian stroma (Figure 3h). On the other hand, the group of fish that received a second injection within the interval of 7 days showed no histological changes in their ovaries after 14 days, when compared to controls (Figure 3i). In kidneys, we observed melanin and lipofuscin pigments, renal thrombosis and autophagy with tubular disarray and loss of tubular lumen epithelium, loss of Bowman’s capsule space and the integrity of the glomerular tuft compromising blood filtration (Figure 3n,o). The frequency of the relative systemic alterations is summarized in Table 1.

**Table 1.**
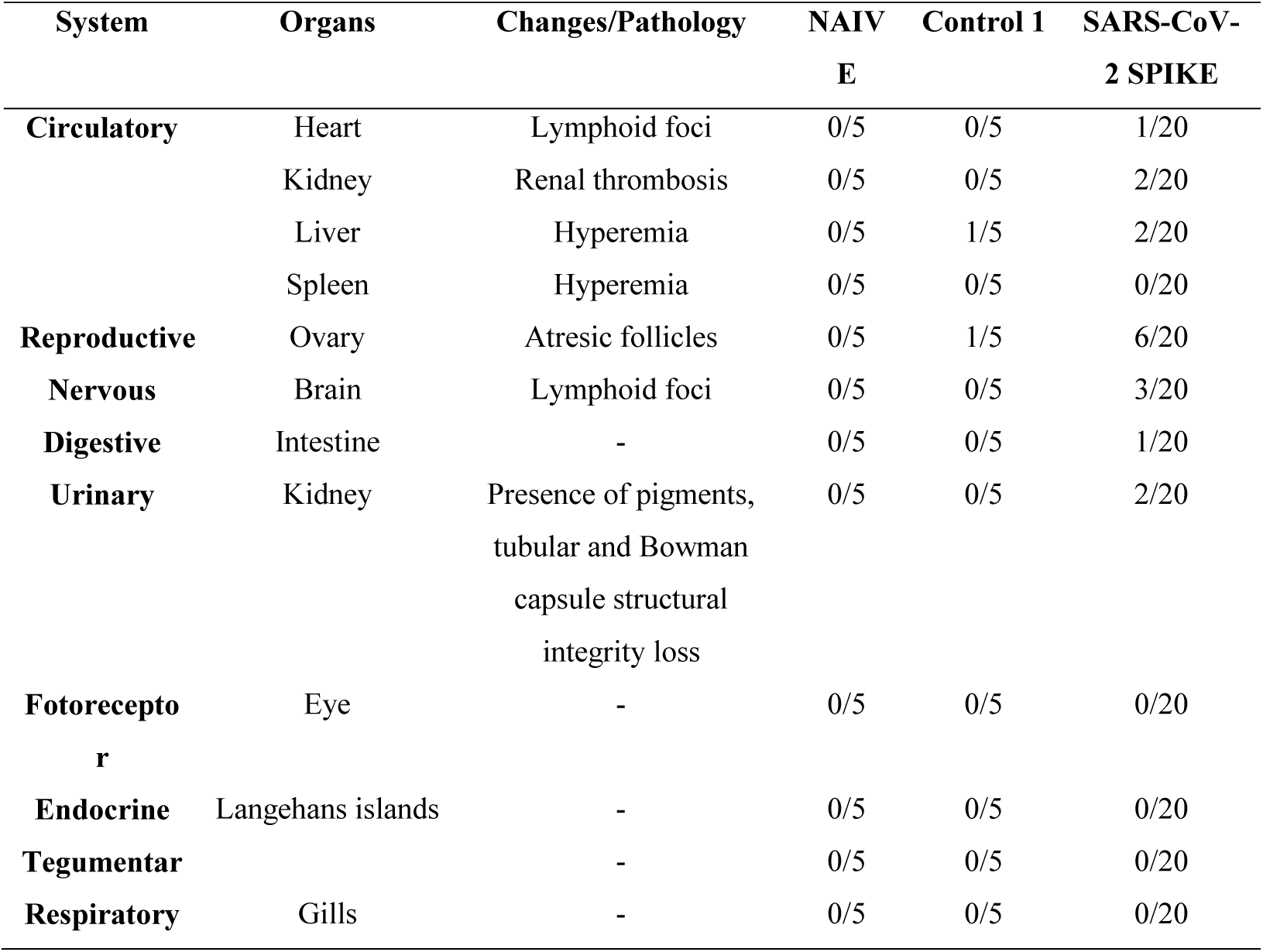
Summary of histopathological findings in different organs of zebrafish injected with rSpike. Number of female fish with histopathological alterations out of total female fish injected. Females were injected either with Naïve control (n = 5), Control 1 (protein buffer) (n = 5), or SARS-CoV-2 protein (n =20).

| System | Organs | Changes/Pathology | NAIV<br>E | Control 1 | SARS-CoV-2 SPIKE |
| --- | --- | --- | --- | --- | --- |
| **Circulatory** | Heart | Lymphoid foci | 0/5 | 0/5 | 1/20 |
|  | Kidney | Renal thrombosis | 0/5 | 0/5 | 2/20 |
|  | Liver | Hyperemia | 0/5 | 1/5 | 2/20 |
|  | Spleen | Hyperemia | 0/5 | 0/5 | 0/20 |
| **Reproductive** | Ovary | Atresic follicles | 0/5 | 1/5 | 6/20 |
| **Nervous** | Brain | Lymphoid foci | 0/5 | 0/5 | 3/20 |
| **Digestive** | Intestine | - | 0/5 | 0/5 | 1/20 |
| **Urinary** | Kidney | Presence of pigments,<br>tubular and Bowman<br>capsule structural<br>integrity loss | 0/5 | 0/5 | 2/20 |
| **Fotoreceptor** | Eye | - | 0/5 | 0/5 | 0/20 |
| **Endocrine** | Langehans islands | - | 0/5 | 0/5 | 0/20 |
| **Tegumentar** |  | - | 0/5 | 0/5 | 0/20 |
| **Respiratory** | Gills | - | 0/5 | 0/5 | 0/20 |

**Figure 3.**
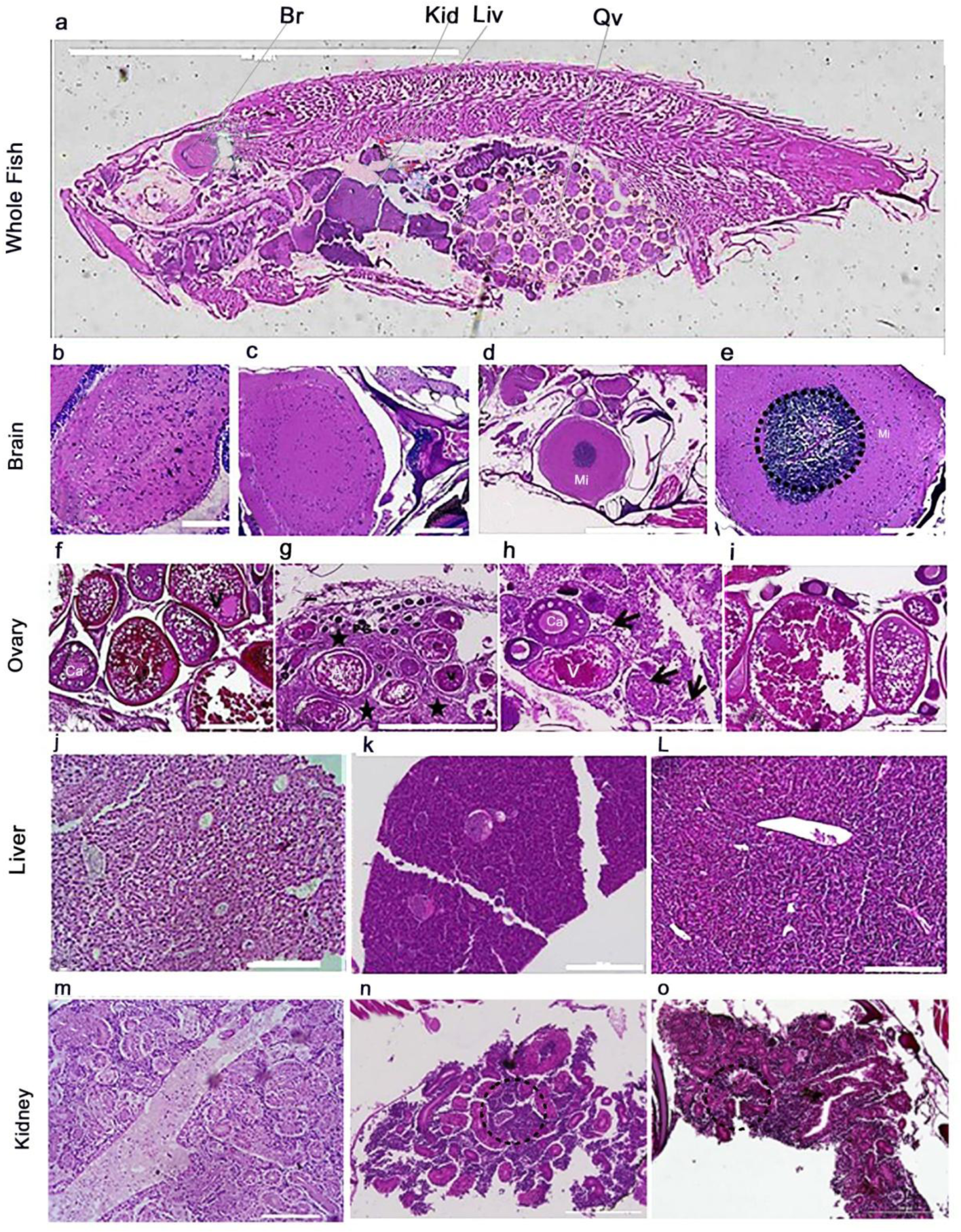
Inflammatory infiltrates in different systems of zebrafish injected with rSpike protein. **a:** longitudinal section of the whole female zebrafish for morphological analyses of the main organs affected. All sections were stained with Hematoxylin Eosin. **Brain:** (b)-histology of control, (c)-brain histology after 7 days of first immunization presenting macrophages, and (d) 14 days after first immunization with a burst after 7 days from the first immunization presenting intense mononuclear infiltrate. (e) The same image as panel d but at a higher magnification. **Ovary:** Ovarian histology from zebrafish control (f), after 7 (g - h) and 14 days (i). (f-i) Follicular development was classified as primary growth oocyte (PG), cortical alveolus (CA), and vitellogenic (V) stages. Asterisks in panel g indicate an abundant and disorganized extracellular matrix in the ovarian stroma. (h) Inset shows a higher magnification of the cellular infiltration and arrows show dense, eosinophilic inflammatory infiltrates. (i) The histology of ovaries after 14 days is similar to the control. Scale bars: 1000 μm (g) and 200 μm (f, h, and i). **Liver:** Histology of the liver from control (j), after 7 days from rSpike immunization (l), and after 14 days from the first immunization with a burst at 7 days (m). **Kidney:** Histology of kidney from zebrafish control (n), after 7 days from the first immunization (o), and after 14 days from the first immunization with a second immunization after 7 days (p). Scale bars: 1,000 μm (n) and 200 μm (o - p).

### rSpike protein immunization induces systemic neutrophils and macrophage infiltration in zebrafish

Taking together clinical evidences of the immunological effects of rSpike protein and the inflammatory-related alterations in the architecture of treated zebrafish tissue, we then turn to a more detailed investigation of the activation of the immune system upon injection of rSpike protein in zebrafish. The presence of the major inflammatory cells as neutrophils and activated macrophages present in the brain and coelomic cavity of the zebrafish were detected by immunostaining. Antibodies against Lymphocyte antigen 6 complex locus G6D (Ly6G), and Allograft inflammatory factor 1 (AIF-1/Iba1) were used to identify neutrophils and activated macrophages, respectively. In non-immunized fish (control group), there was no visible staining for AIF-1/Iba1 (Figure 4 - panel a3); but there was weak staining for Ly6G in the nervous system and ventral area of the coelomic cavity (Figure 4 panel a4). However, the females injected with rSpike protein presented strong Ly6G and AIF-1/Iba1 staining, indicating an inflammatory response of the organism to the virus protein (Figure 4b). Colocalization between Ly6G and AIF-1/Iba1 was observed with predominance in the peripheral region of the brain and in the portion of the kidney from the head (Figure 4b, field 1). Macrophages and neutrophils were also labeled in large vessels (Figure 4b, field 2). In the coelomic cavity in general, there was an increase of neutrophil (Ly6G) and macrophage (AIF/Iba1) cell infiltration (Figure 4c).

**Figure 4.**
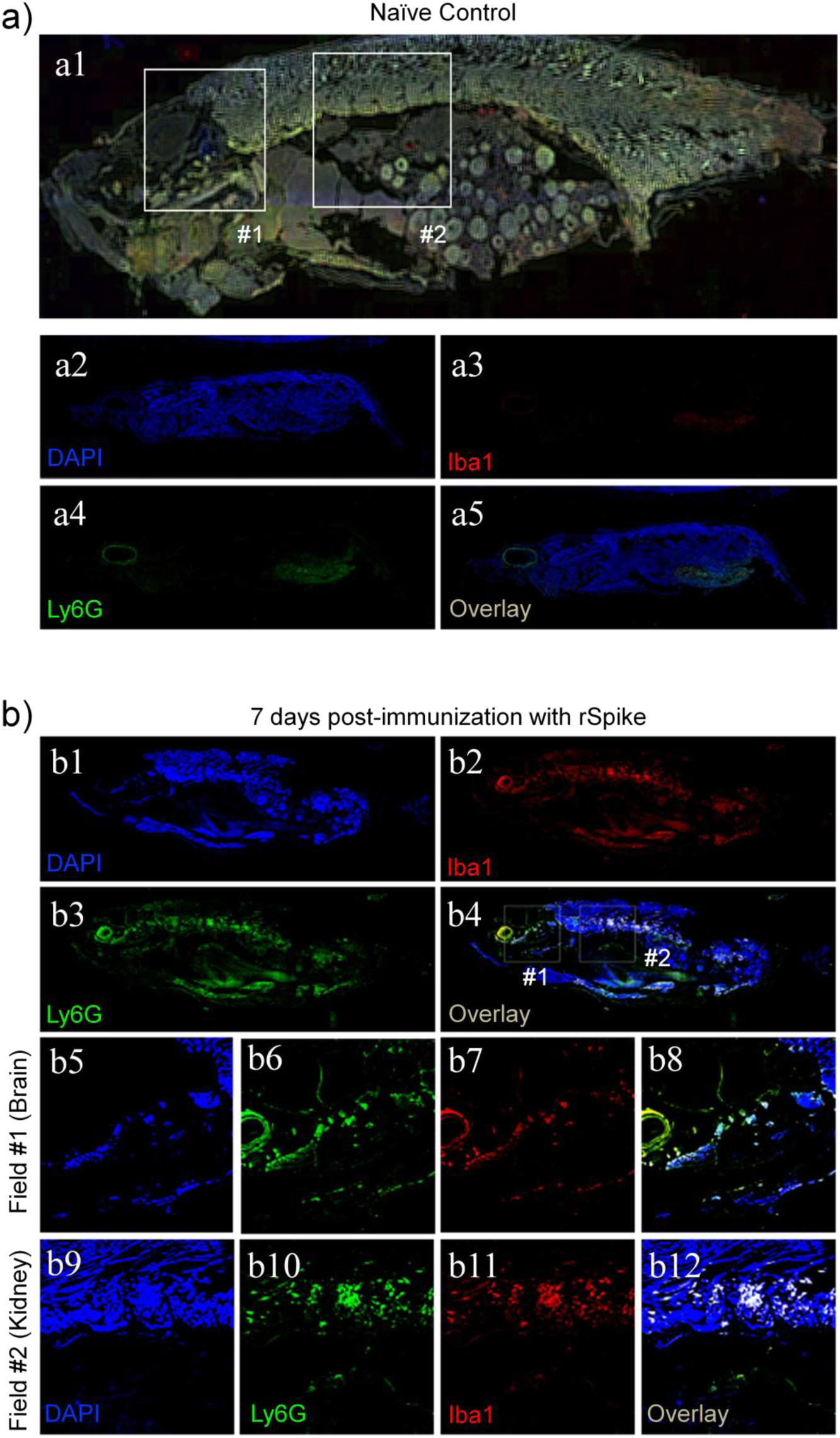
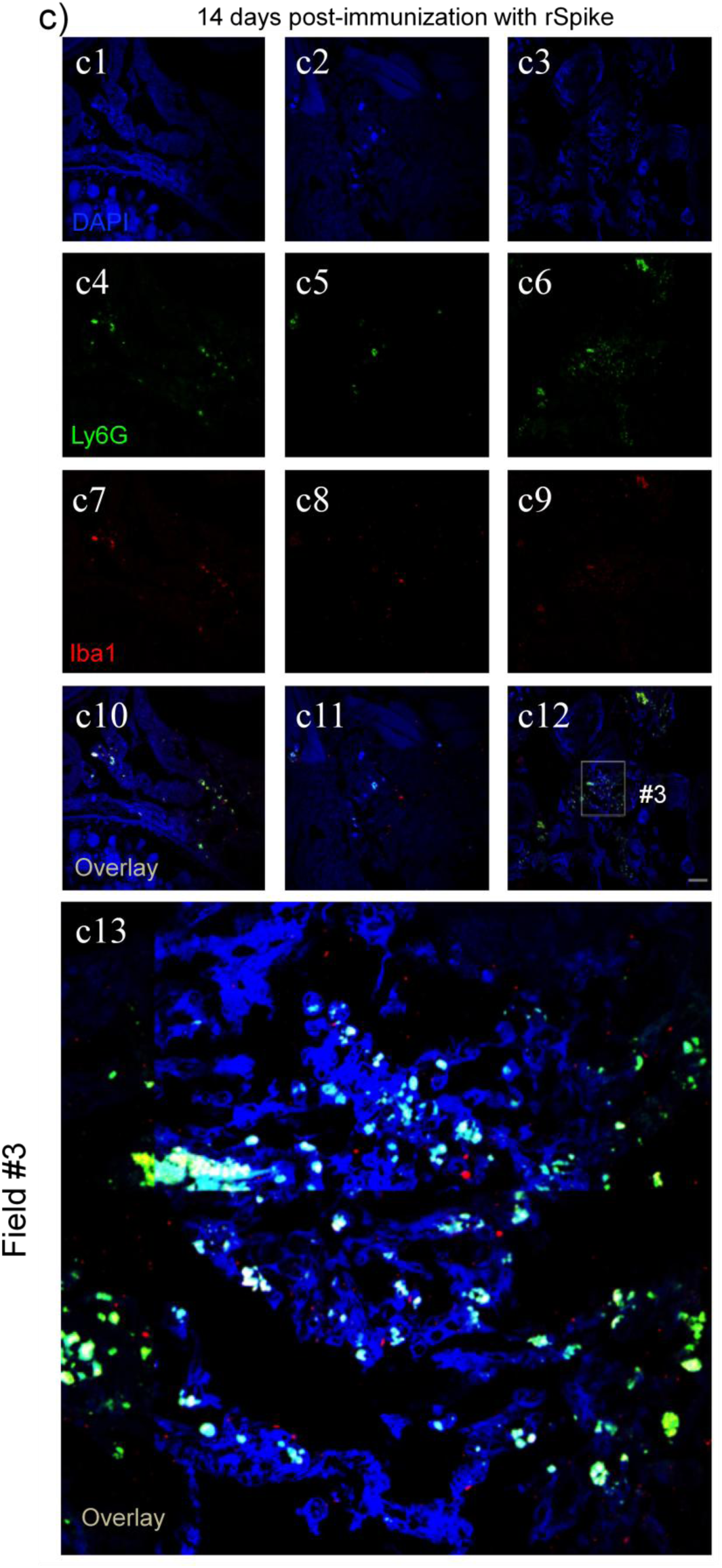
rSpike protein immunization induces systemic neutrophil and macrophage infiltration in zebrafish. Representative immunofluorescence from zebrafish non-immunized control (a) and i.p. immunized with rSpike protein 7 days post-immunization and assessed by scan scope (b). Overview of the whole fish (a1-a5, b1-b4). (c) Confocal-multiphoton imaging from zebrafish immunized twice with rSpike protein after 14 days from the first immunization. The second immunization happened 7 days after the first one. The images depict DAPI (nucleic acid colored in blue), Ly6G (neutrophils colored in green), and Iba1 (macrophages colored in red). Colocalization of DAPI, Ly6G, and Iba1 between fishes are shown in panels described as Overlay. The assay was performed using 7 adult female fishes for immunized groups and adult female fishes for non-immunized group, used as a control group.

The innate immune system and antibody production were detected after rSpike injection in adult zebrafish. The question remained as to whether the fish could respond through cellular immunity, especially T cells. To answer this question, immunostaining revealed the presence of CD-4 and CD-8 cells in the coelomic cavity of female adult zebrafish injected with rSpike (Figure 5).

**Figure 5.**
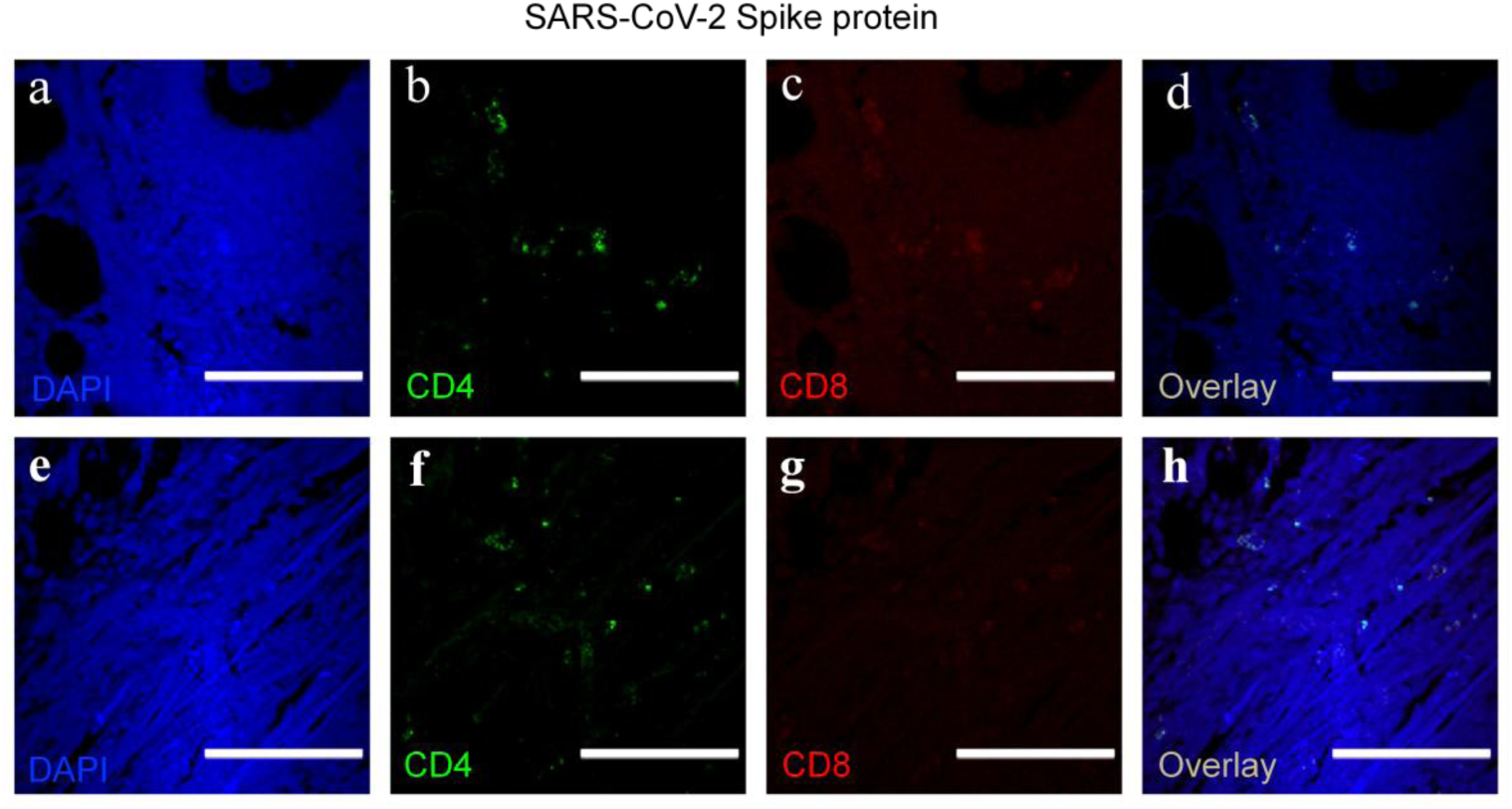
rSpike protein immunization induces innate immune system responses in zebrafish. Immunofluorescence from zebrafish i.p. immunized with rSpike protein after 7 (a-d) and 14 (e-h) days of injection. The images depict DAPI (nucleic acid colored in blue), CD4 (colored in green), and CD8 (colored in red). Colocalization of DAPI, CD4, and CD8 between fish injected are shown in panels d and h (described as Overlay). The assay was performed using 7 adult female fishes for immunized groups.

### The human receptor Angiotensin converting enzyme 2 (ACE2) share 72% sequence similarity to its ortholog in zebrafish

One of the known targets of SARS-CoV-2 Spike protein is the Angiotensin receptor converting enzyme 2 (ACE2) in humans. It is considered the main gateway to the virus infection. Considering the effects of rSpike protein on the fishes analyzed in this work, structural and functional similarities between zebrafish and human ACE2 were investigated, using bioinformatic analysis. Interestingly, zebrafish has ACE2 protein that shares 58 and 72 % primary sequence identity and similarity to human ACE2, respectively (Figure 6; Supplemental Figure 2). Human ACE2 interacts to the receptor binding domain (RBD) of SARS-CoV-2 Spike protein mainly by polar and salt bridge interactions. Human ACE2 has 22 residues making part of the protein-protein interaction and most of them are located at the N-terminal region of ACE2. 77% of the human ACE2 residues of the interface are similar in zebrafish ACE2 sequence (Figure 6; Supplemental Figure 2) suggesting that zebrafish may also binds SARS-CoV-2 Spike protein. The tree-dimensional structure of zebrafish ACE2 based on homology model (Figure 6a) shows a high structural similarity with human ACE2. Computational analysis of protein-protein interaction using ACE2 and the RBD of SARS-CoV-2 Spike protein reveals similar values of binding free energy suggesting that zebrafish is susceptible to virus infection (Figure 6c). In our work, we do not expect that rSpike protein interacts with zebrafish ACE2 because rSpike correspond to the N-terminal part of the Spike protein (residues 16-165) that precedes the RBD domain (residues 319-541).

**Figure 6.**
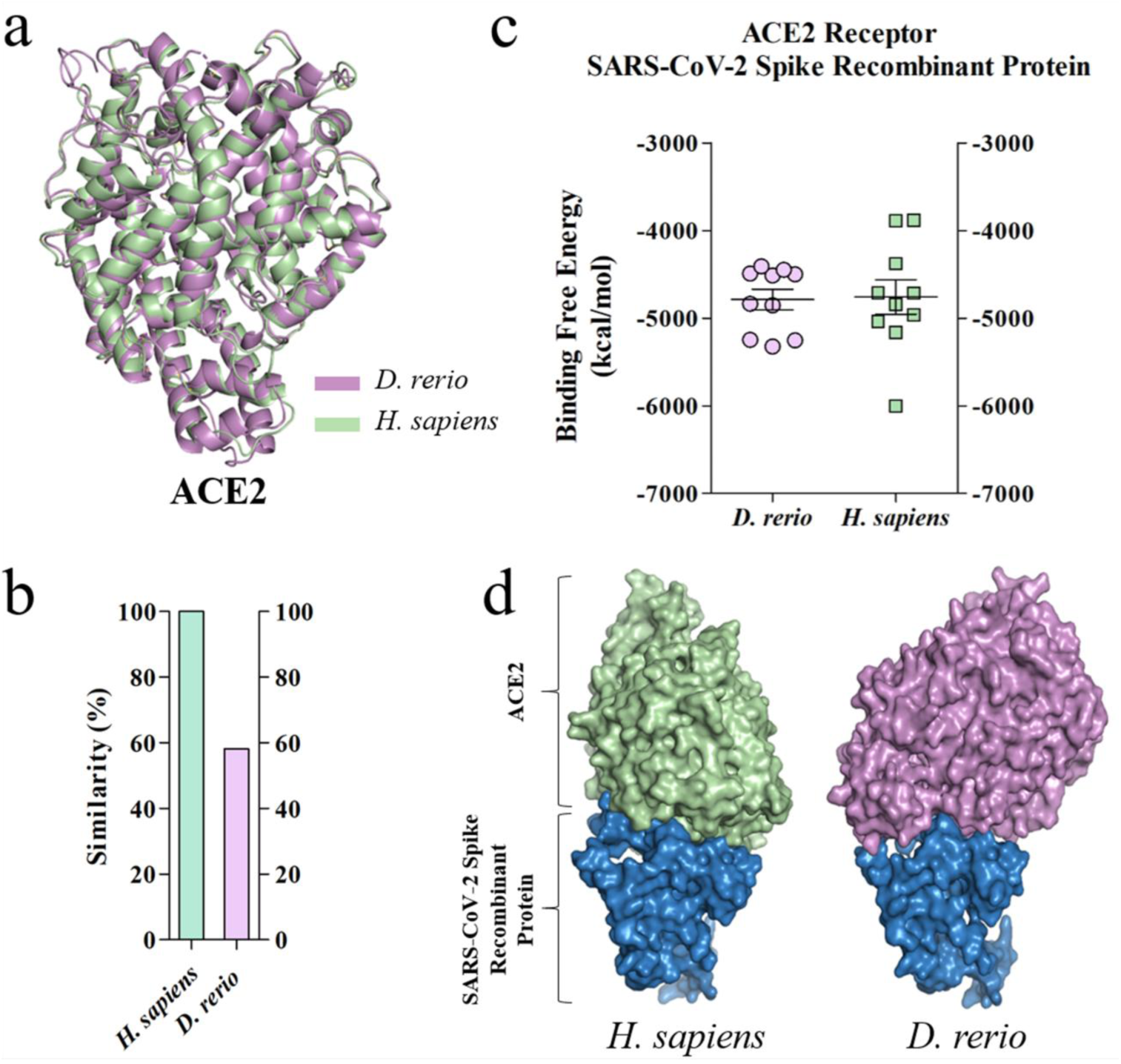
*In silico* analysis of the interaction of the human and zebrafish ACE2 receptor with rSpike protein. (a) Structural alignment between ACE2 of human and zebrafish. For comparison of 3D structures, the FASTA files were converted into PDB files (containing the 3D coordinates of the proteins) using the Raptor X tool (http://raptorx.uchicago.edu). (b) The similarity of ACE2 between human and zebrafish. (c) Graphs show the free binding energy in protein-ligand interactions docking analysis and the axis (X) represents the score of 10 (ten) possibilities of interaction between molecule-ligand and the axis (Y) compares the free binding energy it represents per kilocalorie per mol (Kcal/mol). (Kcal/mol). (d) Protein-protein interaction between human and zebrafish ACE2 and SARS-CoV-2 Spike RBD.

### The protein-protein interaction prediction among SARS-CoV-2

The protein-protein interaction prediction among the rSpike and zebrafish proteins according to the subcellular location (membrane, cytoplasm, and nucleus) predicted interactions with 2,910 proteins for the membrane, 771 proteins for the cytoplasm, and 1,134 proteins for the nucleus (Table 2; Supplementary Table 1). For human proteins and rSpike predicted interactions with 1,785 proteins for the membrane, 1,168 proteins for the cytoplasm, and 1,242 proteins for the nucleus (Table 2; Supplementary Table 1). Considering the most general ontological terms found hierarchically, according to the KEGG and Reactome databases, 71% of the terms identified for zebrafish are identical to those found for human. However, further analysis showed different specific terms with approximately 58% of different specific pathways.

**Table 2.**
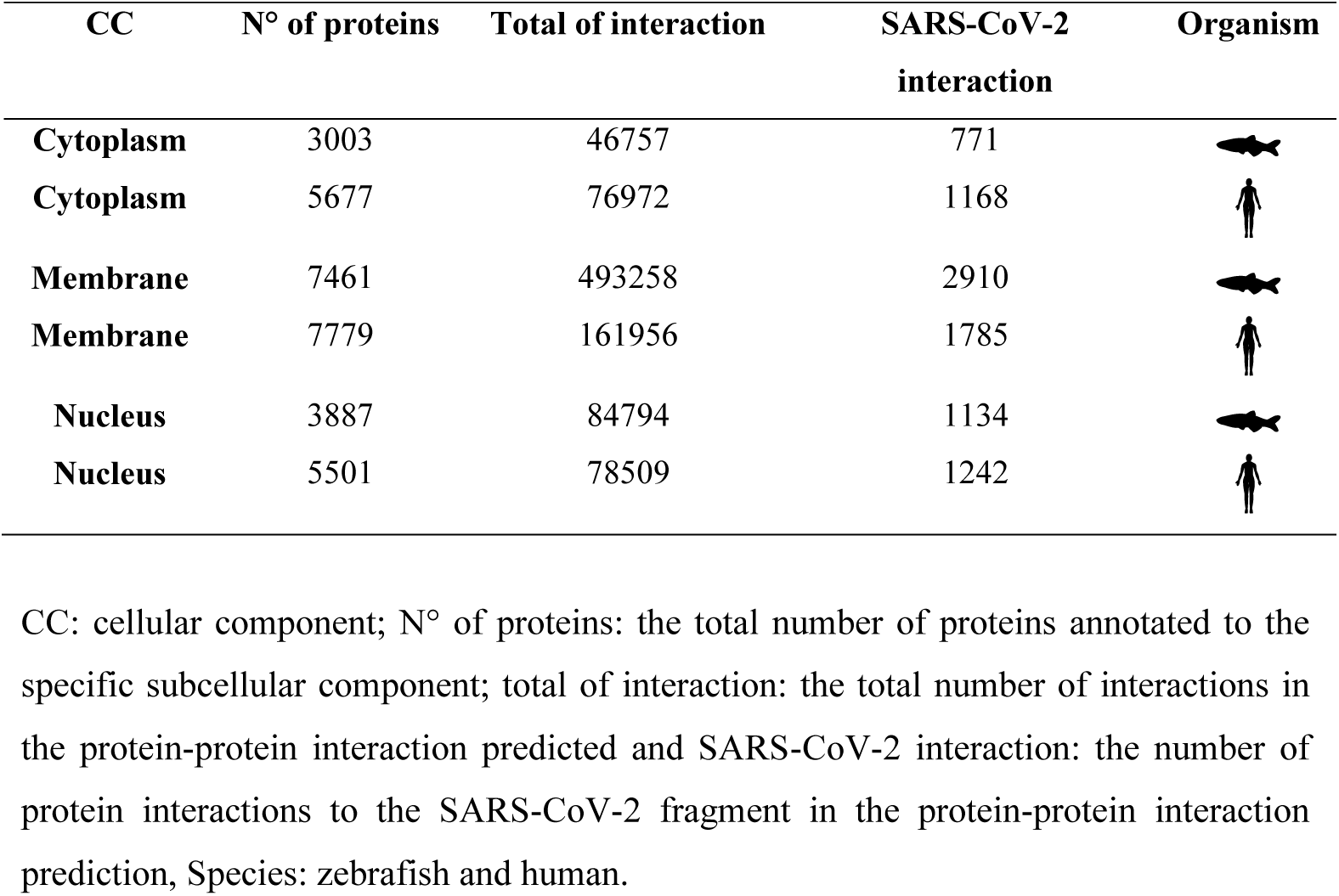
Number of proteins identifying in each cellular component and in the protein-protein interaction prediction.

Functional enrichment of the biological pathways (zebrafish and human) showed basic processes related mainly to cell growth and death, including regulation of transcription and translation mechanisms, mechanisms of DNA repair or replication, and signaling pathways of p53 and by GPCR, among others. Additionally, we identified the pathways related to signal molecules and interactions, signal transduction, and the immune system (Figure 7, Supplementary Table 1).

**Figure 7.**
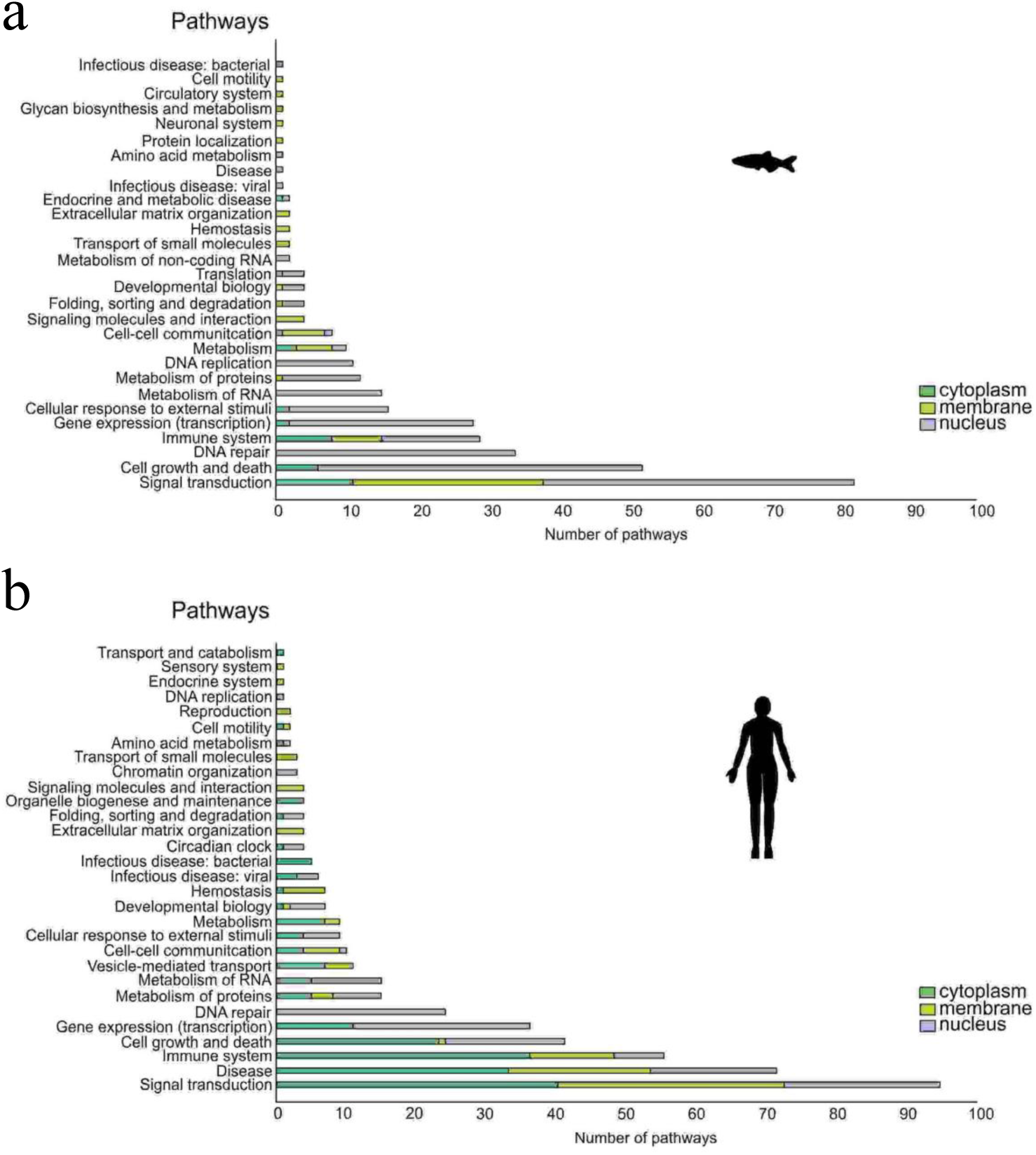
Biological pathways enriched with proteins found from protein-protein interaction prediction with rSpike. Graph relating the proteins from zebrafish (a) and human (b) predicted to interact to rSpike with its cell localization and function within the cell (pathways).

Interestingly, it was recovered through the protein-protein interaction with rSpike, the Toll-like receptor pathway (dre:04620 and hsa:04620). It can allow interaction with the Toll-like receptors TLR1, TLR2, TLR4, and TLR5 and the interferon-α/β receptor (IFNαβR), possibly triggering the activation of various signaling pathways (Figure 8). In this pathway, we observed a possible interaction of the rSpike with the signal transducer and activator of transcription 1-alpha/beta (STAT1) protein in the cytoplasmic region. Additionally, the signal molecules and interaction pathway (zebrafish and human) showed the possibility of rSpike interacting with a considerable number of cell receptors related to the neuroactive ligand receptor (KEGG:4080) and a cytokine-cytokine receptor (Figure 8, KEGG:4060) and triggering diverse cellular signaling such as the TGF beta signaling family, class I and II helical cytokines, IL and TNF family. In addition, proteins related to the extracellular matrix, cellular communication and motility, formation of vesicles, transport and catabolism, VEGF signaling pathway, and AGE-RAGE signaling pathway in diabetic complications, among others, were identified (see Supplementary Table 1).

**Figure 8.**
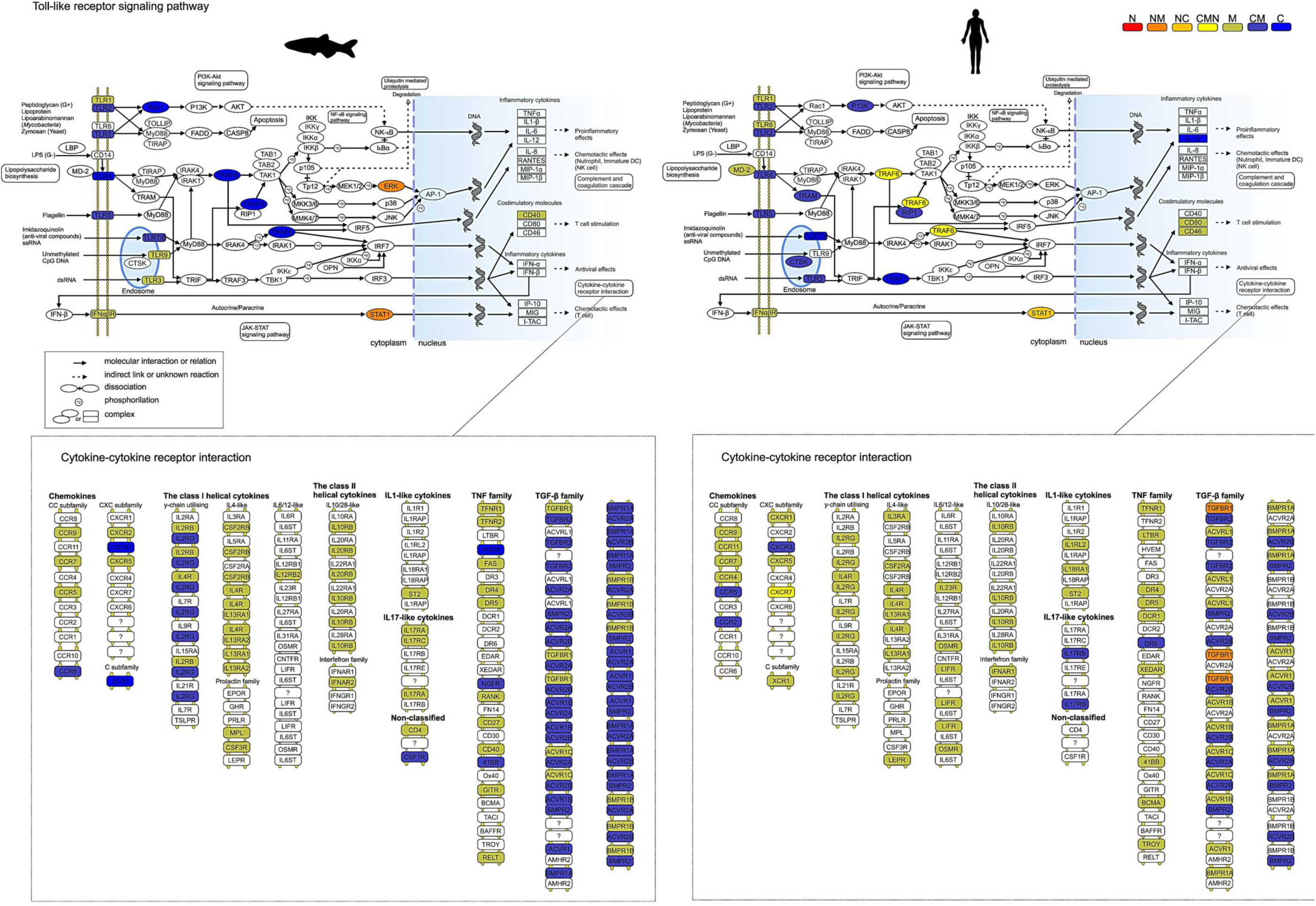
Schematic representation of the Toll-like receptor pathway and cytokine-cytokine receptor interaction. Biological pathway recovered through functional enrichment and mapping of proteins interacting with the recombinant spike protein, rSpike. In red (N) are proteins located in nucleus; dark orange (NM) shows proteins identified in the nucleus and membrane; light orange (NC) shows proteins identified in the nucleus and cytoplasm; yellow (CMN) shows proteins identified in the cytoplasm, membrane, and nucleus; yellow-greenish (M) shows proteins identified in the membrane; dark blue (CM) shows proteins identified in the cytoplasm and membrane; and blue (C) shows proteins identified in the cytoplasm. The schematic represents the zebrafish pathway (a) and the right side of the schematic represents the human pathway (b). The functional enrichment of the pathways was performed with Gprofiler software, and the mapping was performed with the Bioconductor Pathview package. Pathways adapted from KEGG.

The possible virus-host protein interactions during the SARS-CoV-2 infection were tested in network analysis based on protein interactions (Figure 9). The important similarity between SARS-CoV-2 proteome and SARS-CoV proteome^18^ allowed us to hypothesize that the SARS-CoV proteome is highly conserved in SARS-CoV-2. In our network analysis we were able to detect 29 proteins (Figure 9). A PPI interaction database was assembled, including 7 nodes and 29 interactions. We analyzed the following proteins: Parvalbumin 4 (Pvalb4), Creatine kinase (Ckma), Keratin 5 (Krt5), A kinase anchor protein 1 (Ak1), Malate dehydrogenase (Mdh1aa), 2-phospho-D-glycerate hydro-lyase (Eno3), Component Chromosome 15 (ENSDARG00000095050), Component Chromosome 1 (wu:fk65c09), Component Chromosome 16 (Zgc:114037), Component Chromosome 17 9 Zgc:114046), Component Chromosome 26 (ENSDARG00000088889), Apolipoprotein A-II (Apoa2), Apolipoprotein A-Ib (Apoa1b), Serpin peptidase inhibitor member 7 (Serpina7), Transmembrane serine protease 2 (tmprss2), Fetuin B (fetub), Apolipoprotein A-I (apoa1a), Carboxylic ester hydrolase (ces3), Apolipoprotein Bb (apobb), tandem duplicate 1, Fibrinopeptide A (fga), Serotransferrin (tfa), Apolipoprotein C-I (apoc1), Complement component C9 (c9), Pentaxin (crp), Ceruloplasmin (cp), Hemopexin (hpx), Ba1 protein (ba1), Component Chromosome 13 (ENSDARG00000), and Component Chromosome 25 (ENSDARG0000008912).

**Figure 9.**
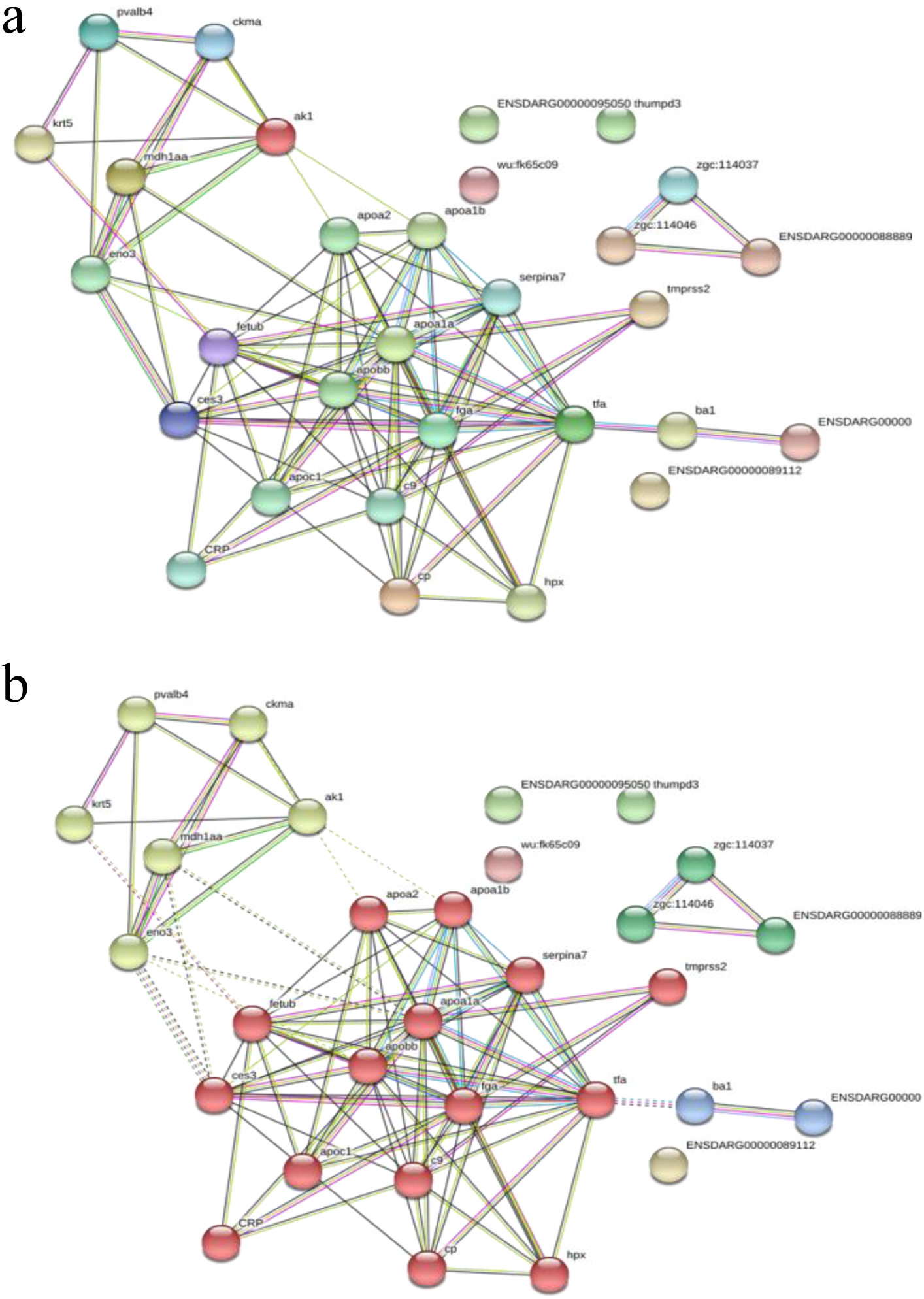
(a and b) Protein interaction network in zebrafish blood plasma. The strongest interactions are exemplified by thicker lines and the weakest are shown by dotted lines. (b) The proteins in red belong to the blood coagulation cascade and also to the immune system pathway. The green proteins are those involved in the structural and chromosome components. The STRING software was used to analyze the protein network and Kyoto Encyclopedia at Genes and Genomes (KEGG) tool to detect protein-protein association. Pvalb4, Parvalbumin 4; Ckma, Creatine kinase; Krt5, Keratin 5; Ak1, A kinase (PRKA) anchor protein 1; Mdh1aa, Malate dehydrogenase; 24 Eno3, 2-phospho-D-glycerate hydro-lyase; ENSDARG00000095050, Component Chromosome 15; wu:fk65c09, Component Chromosome 1; Zgc:114037, Component Chromosome 16; Zgc:114046, Component Chromosome 17; ENSDARG00000088889, Component Chromosome 26; Apoa2, Apolipoprotein A-II; Apoa1b, Apolipoprotein A-Ib; Serpina7, Serpin peptidase inhibitor, clade A (alpha-1 antiproteinase, antitrypsin), member 7; Tmprss2, Transmembrane serine protease 2; Fetub, Fetuin B; Apoa1a, Apolipoprotein A-I; Ces3, Carboxylic ester hydrolase; Apobb, Apolipoprotein Bb, tandem duplicate 1; Fga, Fibrinopeptide A; Tfa, Serotransferrin; Apoc1, Apolipoprotein C-I; C9, Complement component C9; Crp, Pentaxin; Cp, Ceruloplasmin; Hpx, Hemopexin; Ba1, Ba1 protein; ENSDARG00000, Component Chromosome 13; and ENSDARG0000008912, Component Chromosome 25.

## Discussion

Here we show, for the first time, that zebrafish injected with rSpike protein, fragment 16 to 165 (rSpike), that corresponds to the N-terminal portion of the protein, produced an acquired and native immune response and showed adverse effects, following a series of experiments to validate a model of pre-clinical safety studies.

The first experiments aimed to analyze the humoral response with antibody production and used, besides the rSpike, the appropriate negative controls as the *E. coli* extract, mixed of purified recombinant proteins from bacteria, and buffer without the virus protein. There was no increased in computed densitometry of the fragment related to the IgM production in the control groups. Interestingly, the fragment was found only in animals injected with rSpike, demonstrating the specificity of the immune response. Despite the zebrafish systemic antibody production after day 7 of injection of rSpike, the efficiency of these antibodies may have increased on the 14th day, conferring a reduction in the mortality rate of the immunized animals. It was observed using SDS-PAGE that a suggestive time-dependent increase of the fragment correlated with the molecular weight of IgM in zebrafish serum. The results observed in the production of antibodies at 7 and 14 days after inoculation also suggest the similarity to human infected individuals with COVID-19^19^. Antibody production was found in the serum, and in the eggs as well (Figure 1).

In the literature, the passive transfer of antibodies to eggs is known in zebrafish and other teleosts^20^. It has also been described as a strategy for immunization in aquaculture to farmed fish^2^. In evolution, the passive transfer of antibodies protects the offspring from fish to mammals, along with other groups of tetrapods^2,21^. Although in mammals, the IgG is transferred through the placenta and breast milk, in fish the IgM plays this role in the yolk^21^. Similarly, in humans, the presence of SARS-CoV-2 antibodies in breast milk has been found to provide passive immunity for children, to protect them^22,23^. In this sense, the antibody transfer shown in zebrafish could be helpful in future studies to understand the maternal immunological protection of descendants against SARS-CoV-2 or any other vaccine candidates. Also, measuring antibodies against the rSpike in plasma and egg demonstrates great potential for the use of zebrafish in the early stages (phases I and II) of the development and use of these antibodies in therapeutics and prophylactics for humans^24,25^.

Interestingly, fish injected with rSpike produced a toxic inflammatory response with similarity to severe cases of COVID-19 in humans (Figure 2 and 3). Different systems were affected, including the nervous system. The first hint of rSpike toxicity was the distinct swimming behavior that adult females presented after the protein injection. In fact, some recent studies have reported that the SARS-CoV-2 may affect the nervous system^26–28^ as the peripheral nervous system^29–31^, particularly in the most severe cases of infection^32^.

In our study, the rSpike was responsible for generating an inflammatory process in the brain, characterized by an intense influx of mononuclear cells, but no histopathological lesions. This profile is in line with the clinical reports of COVID-19-associated acute necrotizing myelitis^33^, where lymphocytic pleocytosis was observed in the cerebrospinal fluid (CSF). Acute transverse myelitis related to SARS-CoV-2 infection^24^, where an intense leukocyte infiltrate of monocytic characteristic and elevated protein level was also observed in the CSF. In another report, thrombosis in superficial and deep systems, straight sinus, the vein of Galen, internal cerebral veins, and thrombosis of the deep medullary veins were found^27^. Damage to the structure and function of this system can lead to severe encephalitis, toxic encephalopathy, and, after viral infections, severe acute demyelinating lesions^34^. In a case study of 4 children with COVID-19, Abdel-Mannan and collaborators reported that children with COVID-19 may have late neurological symptoms^35^. Future studies with zebrafish might provide more information about the virus damage in the nervous system.

To date, we do not know how the rSpike can cause neurological effects. It is possible that the immune system can recognize these sequence of amino acids. The *in silico* analysis of the rSpike used in the present study indicated that it might interact in a protein-protein level with the Toll-like receptor pathway. In this pathway, the Jak/Stat signaling in humans has been demonstrated to be activated in response to SARS-CoV-2 infection by the release of interleukin IL-6^36,37^. Interestingly, our prediction showed the possible interaction of the rSpike with the signal transducer and activator of transcription 1-alpha/beta (STAT1) protein in the cytoplasmic region, which acts as a carrier for the nucleus and, consequently, executes its function in the inflammatory response as a transcription factor^38–40^. In addition, a study in mice showed that STAT1 deficiency did not affect the response of EGF and other cytokines such as IL10. However, STAT1-deficient mice are more susceptible to pulmonary mycobacterial infection^41^. Similarly, we observed the possibility of interaction with the extracellular signal-regulated kinase 1/2 protein (ERK), in the zebrafish pathway, that plays a role in signaling cascades and produces extracellular signals to intracellular targets^42^.

The response to SARS-CoV-2 in humans appears to be hemophagocytic lymphohistiocytosis (HLH), characterized by immune hyperactivation that occurs when Natural Killer cells and cytotoxic T lymphocytes do not eliminate activated macrophages, leading to excessive production of pro-inflammatory cytokines^43^. These pro-inflammatory cytokines could be associated with a major pathomechanism in kidney damage causing nephrotic proteinuria, collapsing glomerulopathy, membranous glomerulopathy, nephritis, and acute tubular injury^44^. Although some data suggest the incidence of Acute Kidney Injury (AKI) by SARS-CoV-2 to be low^45,46^, other studies indicate that AKI is one of the significantly more common complications in patients who died of COVID-19, pointed out as a marker of multiple organ dysfunction and severe disease^46–48^.

Although we did not measure this cytokine storm, we were able to observe significant renal alterations in the injected animals. In addition, we observed an increase of lymphocyte levels and an increase in melanin and mipofuscin in the kidneys that could be associated with an intense activation of the immune system cells due to the rSpike immunizations response, associated with the accumulation of immune complexes^49^. These results suggest that the immunized fish produced immune complexes.

Histological alterations were analyzed in the liver as mild lobular infiltration by small lymphocytes, centrilobular sinusoidal dilation, patchy necrosis, moderate microvesicular steatosis, mild inflammatory infiltrates in the hepatic lobule, and the portal tract. These changes are similar to those observed in patients with COVID-19^31,48^. Although the zebrafish biochemical liver function was not tested, a three-fold increase in ALT, AST, and GGT levels has been reported during hospitalization^48^. These alterations could be due to the direct cytopathic effect of the virus and could be associated with higher mortality^50^.

With respect to the reproductive tissue, female zebrafish injected with rSpike displayed severe damage in the ovary (follicular atresia, cellular infiltration, and disorganized extracellular matrix) after 7 days of protein inoculation. On the other hand, it is remarkable that ovarian damage was reversed after 14 days, when zebrafish received a second injection of rSpike. In humans, there is evidence that ACE2 mRNA is expressed, at low levels, during all stages of follicle maturation in the ovary^51^, and also in the endometrium^52^. This pattern of ACE2 expression, in line with our observations, could suggest that SARS-CoV-2 affects female fertility in humans and zebrafish. More studies will be necessary to comprehend the molecular mechanisms underlying SARS-CoV-2-induced female infertility and the effects in the ovarian function. To date, damage in the female reproductive system of COVID-19 patients has not been reported yet^53^.

In the sequence of these experimental findings, the *in silico* analysis showed that zebrafish Ace2 receptor has the same potential for protein-ligand interaction as in humans (Figure 6). We show *in silico* and *in vivo* that the zebrafish Ace2 receptor is susceptible to the rSpike and interacts similarly to the human ACE2 receptor. The importance of ACE2 receptor for SARS-CoV-2 infection and its role in vaccine studies is shown in research with transgenic mice (HFH4-hACE2 in C3B6 mice)^54^. The use of ACE2 receptor by SARS-CoV-2 in the attachment and infection of the host cells has been well postulated in mammals, except for murines, and some birds, such as pigeons^55^. The ACE-2 orthologue studies in non-mammalian animals, including zebrafish, suggest the potential to unveil the role of this enzyme and its use for therapeutic purposes^56^.

The receptors associated with the zebrafish humoral and cellular immune response showed structural and functional homology with the human MHC II, MHCI, TCR alpha and beta receptors. Similar results were observed by Bhattacharya and collaborators, who analyzed by docking interaction of 13 peptides with the human MHC I and II receptors and observed the antigenic capacity of these peptides^57^. Our findings provided functional similarity of the same receptors in zebrafish, showing the immunogenic capacity of the alpha and beta TCR receptors, and the functional similarity with the human receptor. The *in silico* data can recognize, process, and present antigens associated with rSpike protein that might be validated in *in vivo* studies in future (Figure 1 a-d). As in mammals, the zebrafish has a conservative adaptive immune system composed of T and B lymphocytes that develop from the thymus and kidneys, respectively. The conservation of the immune system through evolution reveals the importance of fish immunology studies to improve our knowledge of mammalian immunity^58,59^.

The zebrafish enzymatic system is involved in the genetic rearrangement process in which B (BCR) and T lymphocyte receptors (TCR) originate. They also have, like humans, recombinant activating genes that control the gene segments V, D, and J, producing a diversity of antibodies and lymphocyte receptors^60^. Despite this, teleosts produce only three classes of antibodies: IgM^61^, IgW^62^, and IgZ, the latter exclusive to zebrafish^63^. Studies in zebrafish showed that in the regions of the BCR receptor were targets of mutations^5^. Although the affinity of antibodies in ectothermic vertebrates is less efficient than in mammals, the deaminase activation and affinity maturation might contribute to the diversification of antibodies in zebrafish^64,65^.

The world is now experiencing a global campaign to propose and test therapeutics and vaccines. It is imperative to identify animal models for COVID-19 that provide a translational approach for possible successful interventions. From February to October 2020, the findings with animal models for COVID-19 included several candidates, such as mice, Syrian hamsters, ferrets, non-human primates, minks, cats, dogs, pigs, chicken, ducks, and fruit bats^66^. However, no references regarding zebrafish models were found.

Finally, the conserved genetic homology between zebrafish and humans^4^ might be one of the reasons for the intense inflammatory reaction from the immune system of zebrafish to rSpike analyzed in this work. It has provoked damage to organs in a similar pattern as happen in severe cases of COVID-19 in humans. The fish produced innate and acquired immunity that is suitable for future studies to gather valuable information about vaccine responses and therapeutic approaches. Altogether, we present the zebrafish as an animal model for translational COVID-19 research.

## Acknowledgments

Financial and material support was provided through the São Paulo Research Foundation (FAPESP) granted to: Ives Charlie: Fapesp #2018/07098-0; 2019/19939-1; Cristiane Rodrigues Guzzo: Fapesp #2019/00195-2, 2020/04680-0; Chuck Farah: Fapesp #2017/17303-7; Germán G. Sgro: Fapesp #2014/04294-1; Edgar E. Llontop: Fapesp #2019/12234-2; Natalia F. Bueno: Fapesp #2019/18356-2; Camila G. Bomfim: Fapesp #2019/21739-0. LJGB is supported by a research fellowship from Conselho Nacional de Desenvolvimento Científico e Tecnológico, Brazil (CNPq) 303263/2018-0 and FIFG has a PhD fellowship from FAPESP (2019/14285-3). We would like to thank the Medical School Foundation for financial support (Project CG 19,110). We would also like to thank the entire organizing team of the Global Virtual Hackathon 2020 for the award our team received and the support from the Ministry of Transport, Communications and High Technologies of the Republic of Azerbaijan, the United Nations Development Program, and the SUP.VC Acceleration Center. Authors are also thankful to Sartorius for technical support in this work.

## Materials and methods

### Zebrafish maintenance

Wild-type zebrafish from the AB line, and specific pathogen-free (SPF), were raised in Tecniplast Zebtec (Buguggiate, Italy) and maintained in the zebrafish housing systems in the XXXXXXXXXXXXXXX facilities. Fish used for the experiments were obtained from natural crossings and raised according to standard methods^67^. Zebrafish were kept in 3.5 L polycarbonate tanks and fed three times a day with Gemma micro by Skretting (Stavanger, Norway). The photoperiod was 14:10 hours light-dark cycle and the water quality parameters were 28°C ± 2°C; pH = 7.3 ± 0.2; conductivity 500 to 800 µS/cm, referred to as system water. The procedures were approved by the Ethics Committee (CEUA) of the XXXXXXXXXXXXXXXXXXXXXXXXX and registered under protocol number XXXXXXXX.

### Production of recombinant Spike Protein SARS-CoV-2 antigen-based vaccines

Cloning, protein expression, and purification. The DNA fragment coding for the SARS-CoV-2 Spike protein fragment from 16 to 165 (rSpike) was amplified by PCR using SARS-CoV-2 cDNA transcribed from the RNA isolated from the second XXXXXXXXXXX patient, strain HIAE-02:SARS-CoV-2/SP02/human/2020/BRA (GenBank accession number MT126808.1) provided by XXXXXXXXXXXXXXXXXXXXX. The primers used for amplification of the Spike fragment are 5’ AGCATAGCTAGCGTTAATCTTACAACCAGAACTCAATTACC 3’ and 5’ ATTATCGGATCCTTAATTATTCGCACTAGAATAAACTCTGAAC 3’. The PCR product was purified using the GeneJET PCR Purification Kit (Thermo Fisher Scientific) and digested with AnzaTM restriction enzymes *Nhe*I and *BamH*I (Thermo Fisher Scientific). The expression vector used was pET-28a that was also digested with the same pair of restriction enzymes as the amplified rSpike DNA fragment. The digested fragment was used to ligate the rSpike DNA fragment to the digested pET-28a vector using T4 DNA ligase (Thermo Fisher Scientific). The positive clones were confirmed by digestion tests. The rSpike cloned into pET28a results in a protein with a fusion of seven histidine tag at the N-terminal portion of the protein to facilitate the protein purification steps.

rSpike was expressed in *Escherichia coli* strain BL21(DE3) and BL21(DE3) Star. The cells were grown in 2XTY medium (16 g/L of bacto-tryptone, 10g/L of yeast extract, and 5g/L sodium chloride) with added kanamycin (50 µg/ml) under agitation of 200 rpm at 37°C to an OD_600nm_ of 0.6, at which point 0.5 mM isopropyl-β-D-1-thiogalactopyranoside (IPTG) was added. After 4 hours of induction, the cells were collected by centrifugation and stored at 193 K. The cell pellet expressing the rSpike protein was resuspended in lysis buffer (50 mM Tris-HCl pH 7.5, 200 mM NaCl, 5% glycerol, 0.03% Triton-100 and 0.03% Tween-20) and lysed by sonication on an ice bath in a Vibracell VCX750 Ultrasonic Cell Disrupter (Sonics, Newtown, CT, USA). The lysate was centrifuged at 30.000 x g, 4°C for 45 minutes. The pellet fraction was resuspended in 7M urea, 50 mM Tris-HCl pH 7.5, 200 mM NaCl, and 20 mM imidazole on an ice bath under agitation for one hour and centrifuged at 30.000 x g, 4°C for 45 minutes. The soluble fraction was loaded in a HisTrap Chelating HP column (GE Healthcare Life Sciences) previously equilibrated with 7M urea, 50 mM Tris-HCl pH 7.5, 200 mM NaCl, and 20 mM imidazole. Bound proteins were eluted using a linear gradient of imidazole over 20 column volumes (from 20 mM to 1 M imidazole). Fractions with rSpike were concentrated using Amicon Ultra-15 Centrifugal filters (Merck Millipore) with a 3 kDa membrane cutoff and loaded onto a HiLoad 16/600 Superdex 75 pg (GE Healthcare Life Sciences) size exclusion chromatography column previously equilibrated with 7M urea, 50 mM Tris-HCl pH 7.5, 200 mM NaCl, and 1mM EDTA. The eluted fractions containing rSpike protein were analyzed by 15% SDS-PAGE for purity, and the fractions with the target protein were mixed and concentrated using Amicon Ultra-15 Centrifugal filters (Merck Millipore) with a 3 kDa membrane cutoff (Figure 10).

**Figure 10.**
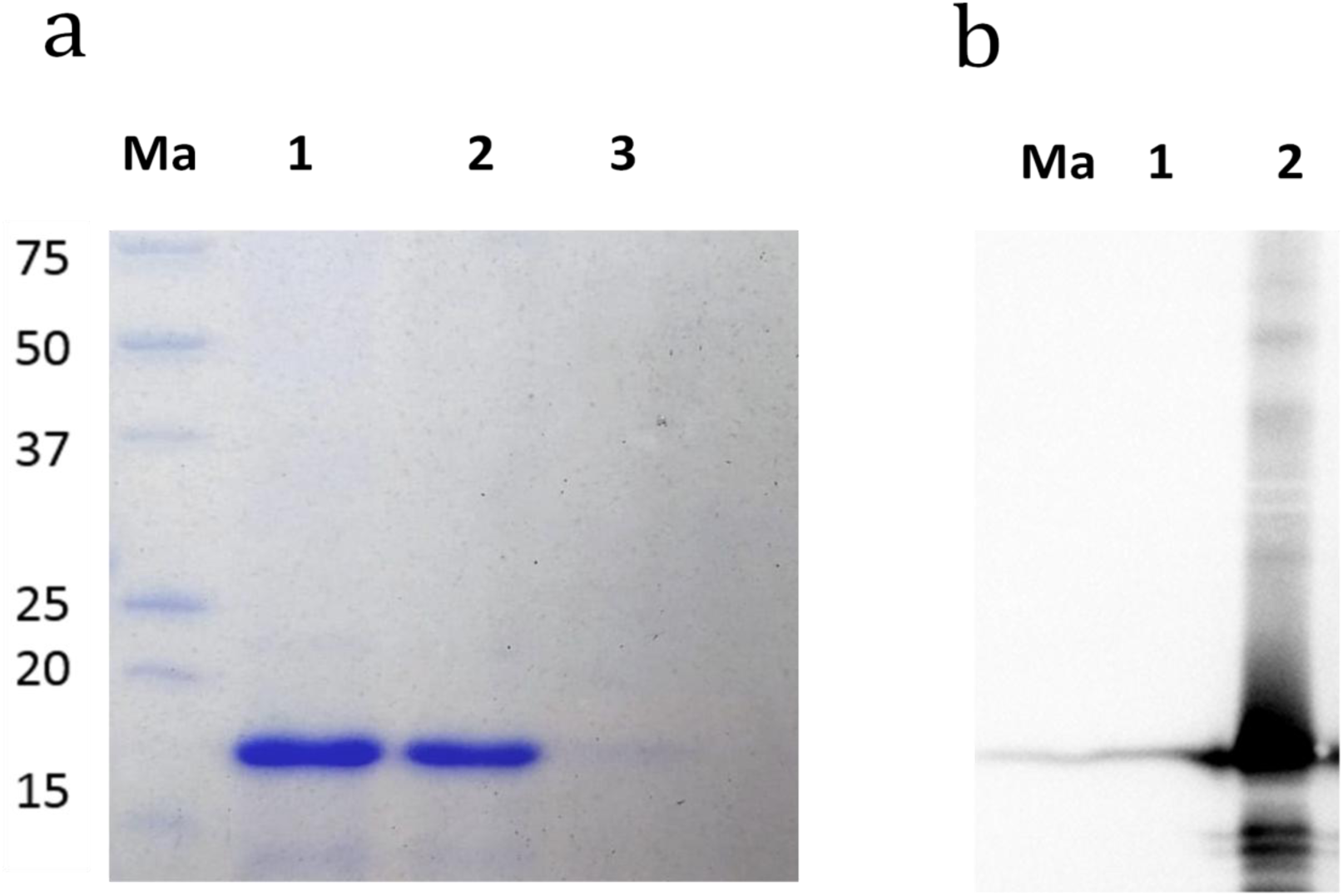
Purification of rSipke protein. a: 15% SDS-PAGE of rSpike purified protein (lanes 1-3) after elution of the protein from exclusion cromatograph column. Ma. Pierce™ Unstained Protein MW Marker (ThermoFisher scientific). b: Western blotting to detect polyhistidine proteins. The protein molecular weight marker (Ma), a not induced *E. coli* BL21(DE3) cells (1) containing the pET28a vector to express rSpike and purified rSpike protein (2) were loaded to a 15% SDS-PAGE and transferred to a nitrocellulose membrane. The membrane was initially blocked with 5% skin milk with PBS buffer for 2 hours and after few PBS rinse the membrane was incubated with monoclonal Anti-polyHistidine−Peroxidase antibody produced in mouse (Sigma-Aldrich).

### The immunization administration

We performed 2 intraperitoneal (IP) inoculations of a solution containing 1 μg purified rSpike diluted in 10 μL of inoculation buffer (7M urea, 50 mM Tris-HCl pH 7.5, 200 mM NaCl, and 1mM EDTA). A group of control animals received injections containing only the dilution buffer. Another control group was challenged by a lysate of bacterial fragment of *E. coli* BL21(DE3) extract. rSpike was injected into two immunization sections in 20 zebrafish females (previously anesthetized with tricaine methanesulfonate (Sigma) - at a dose of 150 mg/L) at an interval of 7 days, with the aim of producing plasma antibodies. Passive antibody transfer to zebrafish eggs occurs naturally as described by Wang and collaborators^20^. After immunization, females were stimulated to mate (at 7 and 14 days after injection) and generated eggs. The time at which the antibodies were transferred to the eggs was analyzed by the western blot technique. Another control group was performed using 1 μg of a mix of proteins in buffer 50 mM Tris-HCl pH 8.0, 200mM NaCl, and 1mM EDTA: equivalent amount of purified PilZ protein from *Xanthomonas citri* (DOI: 10.1016/j.jmb.2009.07.065) and LIC_11128 (residues 1-115 cloned into pET28a expression vector a with a fusion of seven histidine tag at the N-terminal portion of the protein) from *Leptospira interrogans*.

### Antibody responses in zebrafish

Using SDS-PAGE protein electrophoresis, protein from fertilized eggs (10 μg/mL) and serum (10 μg/mL) from adult fish content (after 0, 7 and 14 days) were assessed using methods described by Laemmli^68^. The gels were subsequently stained with 0.25% Coomassie brilliant blue R (Sigma-Aldrich, St. Louis, MO, USA). Molecular weight and protein fraction levels were determined using readings from a computerized densitometer in R software. To identify the protein content, different markers for molecular weights were used and these ranged from 20 to 200 kDa. Protein bands were excised from the SDS-polyacrylamide, and in-gel trypsin digestion was performed according to Shevchenko et al.^69^ and the identification by mass spectrometry. Proteins were precipitated from plasma samples with 4 volumes of cold acetone and 1 volume of cold methanol. The in-solution trypsin digestion was performed according to Lopes Ferreira and collaborators^70^. Mass spectrometric analysis was done by LC– MS/MS.

### Histology from multiple organs

Fixation and decalcification of the adult zebrafish for histology and immunofluorescence was performed according to Moore et al.^71^. For histopathological analysis, 5-μm-thickness sections were mounted on slides and dewaxed in an oven at 60ºC and hydrated in decreasing solutions of xylol three times, and once in xylol + alcohol, for 10 minutes each, followed by a 100, 90, 80, and 70% alcohol battery and washed with distilled water for five minutes. They were then stained with hematoxylin and eosin for observation of the general cellular structures.

### Immunofluorescence assay and image acquisition

For the immunofluorescence assays, the tissues from zebrafish were obtained 0, 7, and 14 days after intraperitoneal injection of SARS-CoV-2 viral protein. Zebrafish tissue sections (5 μm) mounted onto electrically charged slides to increase adherence were deparaffinized in xylol. The samples underwent three 10-minute baths in xylol (P.A.) and a final bath in ethanol/xylol solution (1:1) for 2 minutes. After being deparaffinized, the samples were subjected to hydration by a sequence of ethanol baths at decreasing concentrations (100%, 95%, 90%, 80%, 70%) for 2 minutes in each one, followed by three washes in distilled water. Once hydrated, antigen retrieval was performed by using a trypsin/phosphate-buffered solution (pH 7.2-7.3) mixture (1:1) at 37ºC for 30 minutes in a laboratory drying oven (Thermo Fisher Scientific). Next, the blockade of unspecific epitopes was achieved by a 60-minute incubation in a solution comprised of 2% bovine serum albumin (Sigma Aldrich) (w/v), 0.3% Triton 100X (v/v), and phosphate-buffered solution (pH 7.2-7.3). After that, the primary antibodies were diluted in the aforementioned solution as follows: anti-Ly6G (1:300, Invitrogen, Clone RB6-8C5, Cat 14-5931-81, host: rabbit), anti-AIF-1/Iba1 (1:300, Novus Biologicals, Cat NB100-1028, host: goat) or (1:300, Abcam, Cat ab5076, host: goat), anti-CD4/FITC-conjugated (1:200, eBioscience, Clone RM4-5, Cat 11-0042-85), and anti-CD8/APC.Cy7-conjugated (1:200, BD Bioscience, Clone 53-6.7, Cat 557654). The samples were incubated in these primary antibodies overnight at 4ºC. Then, the samples were washed three times in phosphate-buffered solution (pH 7.2-7.3) for 5 minutes each. Secondary antibodies for anti-Ly6G and anti-Iba1 primary antibodies were diluted as described above, as follows: anti-rabbit/Alexa488 (1:600, Invitrogen, Cat A21206, host: donkey) and anti-goat/Alexa594 (1:600, Invitrogen, Cat A11058, host: donkey). Incubation in these antibodies lasted 2 hours at room temperature. After incubation, the samples were washed three times in phosphate-buffered solution (pH 7.2-7.3) for 10 minutes each. After the final wash, the samples were mounted with a fluoromount containing DAPI dye (VectaShield). Finally, the slides were analyzed under an Olympus VS120 microscope under 20x magnification to acquire images of the whole zebrafish organism before focal analyses of the profile of the immune cells were performed in specific zebrafish structures, which in turn was done by using an Axio Observer combined with LSM 780 confocal device (Carl Zeiss) under the 630x magnification lens.

### Bioinformatics *in silico* analysis

#### Annotation of ontological

The zebrafish and human proteins related to the subcellular location (cytoplasm, membrane, and nucleus) were recovered according to the annotation of ontological terms in the ENSEMBL database (https://www.ensembl.org/index.html, accessed 06/04/2020). For each subcellular location, protein-protein interactions were predicted with a SARS-CoV-2 Spike N-terminal fragment, residues 16-165, (rSpike) using the UNISPPI predictor, where only interactions with a score greater than 0.95 were accepted as interactions^72^. The interacted proteins were submitted to functional enrichment to identify biological pathways using the G:Profiler software^73^, based on the database of zebrafish and human. In addition, the proteins were analyzed with the Bioconductor Pathview package^74^ in the R environment in search of the biological pathways. The pathways were obtained from the Kyoto Encyclopedia of Genes and Genomes (KEGG) database^75^ and the model organism selected was the zebrafish and human.

#### Network analysis

Samples were analyzed in triplicate, and their molecular masses and isoelectric points of the proteins identified by MS / MS were observed using the ProtParam tool (http://us.expasy.org/tools/protparam.html). Data normalization was performed, and a significance cutoff was applied for the identified proteins at log-fold change ± 1.0. Subsequently, the identified proteins on the UniprotKB database were blasted against zebrafish All data obtained were mapped using STRING web tool v11.0 (https://string-db.org/) to screen for protein-protein interactions (PPI).

#### *In silico* analysis

For *in silico* analysis, all FASTA sequences of proteins from zebrafish and human, and SARS-CoV-2 were downloaded from the UNIPROT database (http://www.uniprot.org). We then evaluated the subcellular localization of the identified proteins using the CELLO (subcellular localization predictor) platform v.2.5 (http://cello.life.nctu.edu.tw/) and visualized the proteoform in the cleavage proteins in Protter v. 1.0 (http://wlab.ethz.ch/protter/start/). In addition, the percentage of similarity between the orthologous proteins of different species was calculated using the EMBOSS Water platform (https://www.ebi.ac.uk), and protein alignments were performed using the ESPript platform (http://espript.ibcp.fr/ESPript/cgi-bin/ESPript.cgi). For comparison of 3D structures, the FASTA files were converted into PDB files (containing the 3D coordinates of the proteins) using the Raptor X tool (http://raptorx.uchicago.edu). Then, structural similarities were compared on the iPDA platform (http://www.dsimb.inserm.fr), and structural images of proteins were done using the PyMOL software (https://pymol.org/2/). For the study of protein-protein interaction and Docking of Spike peptides were performed using the Molsoft MolBrower 3.9-1b software.

## Bibliography

1. Peiris M, Leung GM. What can we expect from first-generation COVID-19 vaccines? Lancet. September 2020. doi:10.1016/S0140-6736(20)31976-0

2. Rajan B, Løkka G, Koppang EO, Austbø L. Passive Immunization of Farmed Fish. J Immunol. 2017;198(11):4195–4202. doi:10.4049/jimmunol.1700154

3. Berghmans S, Jette C, Langenau D, et al. Making waves in cancer research: new models in the zebrafish. Biotechniques. 2005;39(2):227–237. doi:10.2144/05392RV02

4. Howe K, Clark MD, Torroja CF, et al. The zebrafish reference genome sequence and its relationship to the human genome. Nature. 2013;496(7446):498–503. doi:10.1038/nature12111

5. Bailone RL, Fukushima HCS, Ventura Fernandes BH, et al. Zebrafish as an alternative animal model in human and animal vaccination research. Lab Anim Res. 2020;36(1):13. doi:10.1186/s42826-020-00042-4

6. Charlie-Silva I, Feitosa NM, Gomes JMM, et al. Potential of mucoadhesive nanocapsules in drug release and toxicology in zebrafish. Mukherjee A, ed. PLoS One. 2020;15(9):e0238823. doi:10.1371/journal.pone.0238823

7. MacRae CA, Peterson RT. Zebrafish as tools for drug discovery. Nat Rev Drug Discov. 2015;14(10):721–731. doi:10.1038/nrd4627

8. Lurie N, Saville M, Hatchett R, Halton J. Developing Covid-19 Vaccines at Pandemic Speed. N Engl J Med. 2020;382(21):1969–1973. doi:10.1056/NEJMp2005630

9. Singh A, Singh RS, Sarma P, et al. A Comprehensive Review of Animal Models for Coronaviruses: SARS-CoV-2, SARS-CoV, and MERS-CoV. Virol Sin. 2020;35(3):290–304. doi:10.1007/s12250-020-00252-z

10. Lu S, Zhao Y, Yu W, Yang Y, Gao J, Wang J. Comparison of SARS-CoV-2 infections among 3 species of non-human Affiliations: 1–27.

11. Niu J, Shen L, Huang B, et al. Non-invasive bioluminescence imaging of HCoV-OC43 infection and therapy in the central nervous system of live mice. Antiviral Res. 2020;173:104646. doi:10.1016/j.antiviral.2019.104646

12. Hanke L, Vidakovics Perez L, Sheward DJ, et al. An alpaca nanobody neutralizes SARS-CoV-2 by blocking receptor interaction. Nat Commun. 2020;11(1):1–9. doi:10.1038/s41467-020-18174-5

13. Cunha LE, Stolet A, Strauch M, et al. Equine hyperimmune globulin raised against the SARS-CoV-2 spike glycoprotein has extremely high neutralizing titers. 2020:1–28. doi:https://doi.org/10.1101/2020.08.17.254375.

14. Zylberman V, Sanguineti S, Pontoriero A V, et al. Development of a hyperimmune equine serum therapy for COVID-19 in Argentina. Medicina (B Aires). 2020;80 Suppl 3:1–6. http://www.ncbi.nlm.nih.gov/pubmed/32658841.

15. Deng J, Jin Y, Liu Y, et al. Serological survey of SARS-CoV-2 for experimental, domestic, companion and wild animals excludes intermediate hosts of 35 different species of animals. Transbound Emerg Dis. 2020;67(4):1745–1749. doi:10.1111/tbed.13577

16. Galindo-Villegas J. The Zebrafish Disease and Drug Screening Model: A Strong Ally Against Covid-19. Front Pharmacol. 2020;11. doi:10.3389/fphar.2020.00680

17. Gaudin R, Goetz JG. Tracking Mechanisms of Viral Dissemination In Vivo. Trends Cell Biol. October 2020. doi:10.1016/j.tcb.2020.09.005

18. Sawicki SG, Sawicki DL, Siddell SG. A Contemporary View of Coronavirus Transcription. J Virol. 2007;81(1):20–29. doi:10.1128/JVI.01358-06

19. Long QX, Liu BZ, Deng HJ, et al. Antibody responses to SARS-CoV-2 in patients with COVID-19. Nat Med. 2020;26(6):845–848. doi:10.1038/s41591-020-0897-1

20. Wang H, Ji D, Shao J, Zhang S. Maternal transfer and protective role of antibodies in zebrafish Danio rerio. Mol Immunol. 2012;51(3-4):332–336. doi:10.1016/j.molimm.2012.04.003

21. Swain P, Nayak SK. Role of maternally derived immunity in fish. Fish Shellfish Immunol. 2009;27(2):89–99. doi:10.1016/j.fsi.2009.04.008

22. Yu Y, Xu J, Li Y, Hu Y, Li B. Breast Milk-fed Infant of COVID-19 Pneumonia Mother: a Case Report. 2020:1–11. doi:10.21203/rs.3.rs-20792/v1

23. Demers-Mathieu V, Dung M, Mathijssen GB, et al. Difference in levels of SARS-CoV-2 S1 and S2 subunits- and nucleocapsid protein-reactive SIgM/IgM, IgG and SIgA/IgA antibodies in human milk. J Perinatol. September 2020. doi:10.1038/s41372-020-00805-w

24. Munz M, Wessendorf S, Koretsis G, et al. Acute transverse myelitis after COVID-19 pneumonia. J Neurol. 2020;267(8):2196–2197. doi:10.1007/s00415-020-09934-w

25. Xu Z, Shi L, Wang Y, et al. Pathological findings of COVID-19 associated with acute respiratory distress syndrome. Lancet Respir Med. 2020;8(4):420–422. doi:10.1016/S2213-2600(20)30076-X

26. Lu Y, Li X, Geng D, et al. Cerebral Micro-Structural Changes in COVID-19 Patients – An MRI-based 3-month Follow-up Study. EClinicalMedicine. 2020;25:100484. doi:10.1016/j.eclinm.2020.100484

27. Cavalcanti DD, Raz E, Shapiro M, et al. Cerebral Venous Thrombosis Associated with COVID-19. Am J Neuroradiol. 2020;41(8):1370–1376. doi:10.3174/ajnr.A6644

28. Iadecola C, Anrather J, Kamel H. Effects of COVID-19 on the Nervous System. Cell. August 2020. doi:10.1016/j.cell.2020.08.028

29. Lau K-K, Yu W-C, Chu C-M, Lau S-T, Sheng B, Yuen K-Y. Possible Central Nervous System Infection by SARS Coronavirus. Emerg Infect Dis. 2004;10(2):342–344. doi:10.3201/eid1002.030638

30. Netland J, Meyerholz DK, Moore S, Cassell M, Perlman S. Severe Acute Respiratory Syndrome Coronavirus Infection Causes Neuronal Death in the Absence of Encephalitis in Mice Transgenic for Human ACE2. J Virol. 2008;82(15):7264–7275. doi:10.1128/JVI.00737-08

31. Tian S, Xiong Y, Liu H, et al. Pathological study of the 2019 novel coronavirus disease (COVID-19) through postmortem core biopsies. Mod Pathol. 2020;33(6):1007–1014. doi:10.1038/s41379-020-0536-x

32. Beghi E, Feigin V, Caso V, Santalucia P, Logroscino G. COVID-19 Infection and Neurological Complications: Present Findings and Future Predictions. Neuroepidemiology. July 2020:1–6. doi:10.1159/000508991

33. Sotoca J, Rodríguez-Álvarez Y. COVID-19-associated acute necrotizing myelitis. Neurol - Neuroimmunol Neuroinflammation. 2020;7(5):e803. doi:10.1212/NXI.0000000000000803

34. Wright EJ, Brew BJ, Wesselingh SL. Pathogenesis and Diagnosis of Viral Infections of the Nervous System. Neurol Clin. 2008;26(3):617–633. doi:10.1016/j.ncl.2008.03.006

35. Abdel-Mannan O, Eyre M, Löbel U, et al. Neurologic and Radiographic Findings Associated With COVID-19 Infection in Children. JAMA Neurol. July 2020. doi:10.1001/jamaneurol.2020.2687

36. Campochiaro C, Dagna L. The conundrum of interleukin-6 blockade in COVID-19. Lancet Rheumatol. 2020;2(10):e579–e580. doi:10.1016/S2665-9913(20)30287-3

37. Sinha P, Mostaghim A, Bielick CG, et al. Early administration of interleukin-6 inhibitors for patients with severe COVID-19 disease is associated with decreased intubation, reduced mortality, and increased discharge. Int J Infect Dis. 2020;99:28–33. doi:10.1016/j.ijid.2020.07.023

38. Casanova J-L, Holland SM, Notarangelo LD. Inborn Errors of Human JAKs and STATs. Immunity. 2012;36(4):515–528. doi:10.1016/j.immuni.2012.03.016

39. Baris S, Alroqi F, Kiykim A, et al. Severe Early-Onset Combined Immunodeficiency due to Heterozygous Gain-of-Function Mutations in STAT1. J Clin Immunol. 2016;36(7):641–648. doi:10.1007/s10875-016-0312-3

40. Goswami R, Kaplan MH. STAT Transcription Factors in T Cell Control of Health and Disease. In:; 2017:123–180. doi:10.1016/bs.ircmb.2016.09.012

41. Sugawara I, Yamada H, Mizuno S. STAT1 Knockout Mice are Highly Susceptible to Pulmonary Mycobacterial Infection. Tohoku J Exp Med. 2004;202(1):41–50. doi:10.1620/tjem.202.41

42. Guo Y, Pan W, Liu S, Shen Z, Xu Y, Hu L. ERK/MAPK signalling pathway and tumorigenesis (Review). Exp Ther Med. January 2020. doi:10.3892/etm.2020.8454

43. Janka GE, Lehmberg K. Hemophagocytic syndromes — An update. Blood Rev. 2014;28(4):135–142. doi:10.1016/j.blre.2014.03.002

44. Kudose S, Batal I, Santoriello D, et al. Kidney Biopsy Findings in Patients with COVID-19. J Am Soc Nephrol. 2020;31(9):1959–1968. doi:10.1681/ASN.2020060802

45. Pan X, Xu D, Zhang H, Zhou W, Wang L, Cui X. Identification of a potential mechanism of acute kidney injury during the COVID-19 outbreak: a study based on single-cell transcriptome analysis. Intensive Care Med. 2020;46(6):1114–1116. doi:10.1007/s00134-020-06026-1

46. Ronco C, Reis T. Kidney involvement in COVID-19 and rationale for extracorporeal therapies. Nat Rev Nephrol. 2020;16(6):308–310. doi:10.1038/s41581-020-0284-7

47. Deng Y, Liu W, Liu K, et al. Clinical characteristics of fatal and recovered cases of coronavirus disease 2019 in Wuhan, China: a retrospective study. Chin Med J (Engl). 2020;133(11):1261–1267. doi:10.1097/CM9.0000000000000824

48. Cai Q, Huang D, Yu H, et al. COVID-19: Abnormal liver function tests. J Hepatol. 2020;73(3):566–574. doi:10.1016/j.jhep.2020.04.006

49. Agius C, Roberts RJ. Melano-macrophage centres and their role in fish pathology. J Fish Dis. 2003;26(9):499–509. doi:10.1046/j.1365-2761.2003.00485.x

50. Jothimani D, Venugopal R, Abedin MF, Kaliamoorthy I, Rela M. COVID-19 and the liver. J Hepatol. June 2020. doi:10.1016/j.jhep.2020.06.006

51. Reis FM, Bouissou DR, Pereira VM, Camargos AF, dos Reis AM, Santos RA. Angiotensin-(1-7), its receptor Mas, and the angiotensin-converting enzyme type 2 are expressed in the human ovary. Fertil Steril. 2011;95(1):176–181. doi:10.1016/j.fertnstert.2010.06.060

52. Vaz-Silva J, Carneiro MM, Ferreira MC, et al. The Vasoactive Peptide Angiotensin-(1—7), Its Receptor Mas and the Angiotensin-converting Enzyme Type 2 are Expressed in the Human Endometrium. Reprod Sci. 2009;16(3):247–256. doi:10.1177/1933719108327593

53. Zupin L, Pascolo L, Zito G, Ricci G, Crovella S. SARS-CoV-2 and the next generations: which impact on reproductive tissues? J Assist Reprod Genet. August 2020. doi:10.1007/s10815-020-01917-0

54. Jiang R-D, Liu M-Q, Chen Y, et al. Pathogenesis of SARS-CoV-2 in Transgenic Mice Expressing Human Angiotensin-Converting Enzyme 2. Cell. 2020;182(1):50-58.e8. doi:10.1016/j.cell.2020.05.027

55. Qiu Y, Zhao Y-B, Wang Q, et al. Predicting the angiotensin converting enzyme 2 (ACE2) utilizing capability as the receptor of SARS-CoV-2. Microbes Infect. 2020;22(4-5):221–225. doi:10.1016/j.micinf.2020.03.003

56. Chou CF, Loh CB, Foo YK, et al. ACE2 orthologues in non-mammalian vertebrates (Danio, Gallus, Fugu, Tetraodon and Xenopus). Gene. 2006;377(1-2):46–55. doi:10.1016/j.gene.2006.03.010

57. Bhattacharya M, Sharma AR, Patra P, et al. Development of epitope-based peptide vaccine against novel coronavirus 2019 (SARS-COV-2): Immunoinformatics approach. J Med Virol. 2020;92(6):618–631. doi:10.1002/jmv.25736

58. Sunyer JO. Fishing for mammalian paradigms in the teleost immune system. Nat Immunol. 2013;14(4):320–326. doi:10.1038/ni.2549

59. Flajnik MF. A cold-blooded view of adaptive immunity. Nat Rev Immunol. 2018;18(7):438–453. doi:10.1038/s41577-018-0003-9

60. Litman GW, Cannon JP, Dishaw LJ. Reconstructing immune phylogeny: new perspectives. Nat Rev Immunol. 2005;5(11):866–879. doi:10.1038/nri1712

61. Mashoof S, Criscitiello M. Fish Immunoglobulins. Biology (Basel). 2016;5(4):45. doi:10.3390/biology5040045

62. Rumfelt LL, Lohr RL, Dooley H, Flajnik MF. Diversity and repertoire of IgW and IgM VH families in the newborn nurse shark. BMC Immunol. 2004;5:1–15. doi:10.1186/1471-2172-5-8

63. Hu Y-L, Xiang L-X, Shao J-Z. Identification and characterization of a novel immunoglobulin Z isotype in zebrafish: Implications for a distinct B cell receptor in lower vertebrates. Mol Immunol. 2010;47(4):738–746. doi:10.1016/j.molimm.2009.10.010

64. Marianes AE, Zimmerman AM. Targets of somatic hypermutation within immunoglobulin light chain genes in zebrafish. Immunology. 2011;132(2):240–255. doi:10.1111/j.1365-2567.2010.03358.x

65. Lewis KL, Del Cid N, Traver D. Perspectives on antigen presenting cells in zebrafish. Dev Comp Immunol. 2014;46(1):63–73. doi:10.1016/j.dci.2014.03.010

66. Muñoz-Fontela C, Dowling WE, Funnell SGP, et al. Animal models for COVID-19. Nature. 2020. doi:10.1038/s41586-020-2787-6

67. Tsang B, Zahid H, Ansari R, Lee RC-Y, Partap A, Gerlai R. Breeding Zebrafish: A Review of Different Methods and a Discussion on Standardization. Zebrafish. 2017;14(6):561–573. doi:10.1089/zeb.2017.1477

68. Laemmli UK. 227680a0. Nature. 1970;227:680–685.

69. Shevchenko G, Konzer A, Musunuri S, Bergquist J. Neuroproteomics tools in clinical practice. Biochim Biophys Acta - Proteins Proteomics. 2015;1854(7):705–717. doi:10.1016/j.bbapap.2015.01.016

70. Babaei E, Alilu S, Laali S. A New General Topology for Cascaded Multilevel Inverters With Reduced Number of Components Based on Developed H-Bridge. IEEE Trans Ind Electron. 2014;61(8):3932–3939. doi:10.1109/TIE.2013.2286561

71. Moore JL, Aros M, Steudel KG, Cheng KC. Fixation and Decalcification of Adult Zebrafish for Histological, Immunocytochemical, and Genotypic Analysis. Biotechniques. 2002;32(2):296–298. doi:10.2144/02322st03

72. Valente GT, Acencio ML, Martins C, Lemke N. The Development of a Universal In Silico Predictor of Protein-Protein Interactions. Csermely P, ed. PLoS One. 2013;8(5):e65587. doi:10.1371/journal.pone.0065587

73. Reimand J, Arak T, Adler P, et al. g:Profiler—a web server for functional interpretation of gene lists (2016 update). Nucleic Acids Res. 2016;44(W1):W83–W89. doi:10.1093/nar/gkw199

74. Luo W, Brouwer C. Pathview: an R/Bioconductor package for pathway-based data integration and visualization. Bioinformatics. 2013;29(14):1830–1831. doi:10.1093/bioinformatics/btt285

75. Ogata H, Goto S, Sato K, Fujibuchi W, Bono H, Kanehisa M. KEGG: Kyoto Encyclopedia of Genes and Genomes. Nucleic Acids Res. 1999;27(1):29–34. doi:10.1093/nar/27.1.29

