## Supplementary figures and images for "Zebrafish studies on the vaccine candidate to COVID-19, the Spike protein: Production of antibody and adverse reaction"

### Supplemental Figure 1

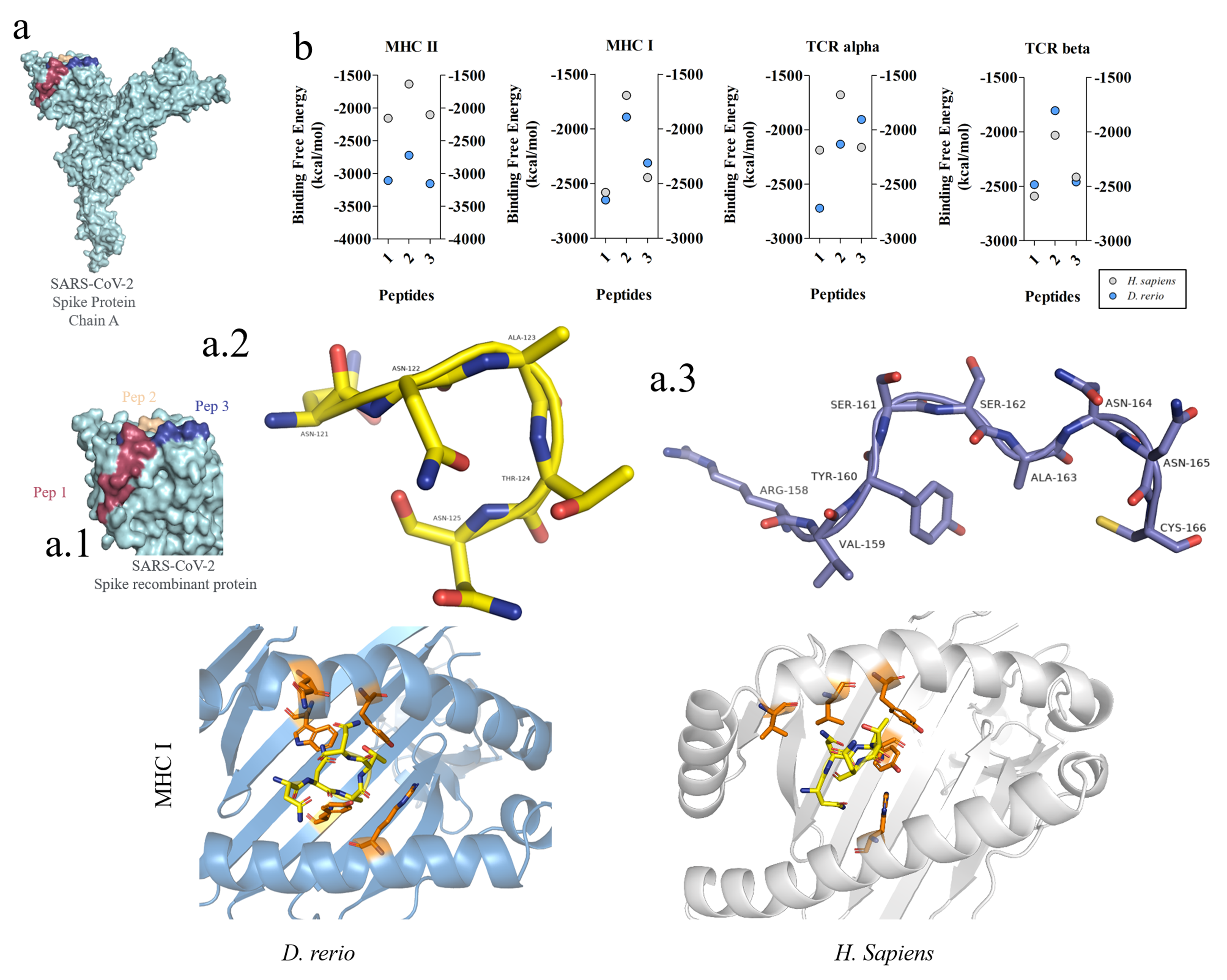
