## Supplemental Figure 2 for "Zebrafish studies on the vaccine candidate to COVID-19, the Spike protein: Production of antibody and adverse reaction"

```
ACE2_HUMAN      MSSSSWLLLSLVAVTAAQSTIEEQAKTFLDKFNHEAEDLFYQSSLASWNYNTNITEENVQ
ACE2_fish       -MCARWLLLLALASVACCQTVEDRAREFLNKFDEEASDIMYQYTLASWAYNTDISQENAD
                  .: ****  :* .*. .*:*::*: **:**:.**.*::** :**** ***:*::**.:

ACE2_HUMAN      NMNNAGDKWSAFLKEQSTLAQMYPLQEIQNLTVKLQLQALQQNGSSVLSEDKSKRLNTIL
ACE2_fish       KEAEAYAIWSEYYNKMSEESNAYPIDQISDPIIKMQLQKLQDKGSGALSPDKASELRNIM
                :  :*   ** : :: *  :: **:::*.:  :*:*** **::**..** **:..*..*:

ACE2_HUMAN      NTMSTIYSTGKVCNPDNPQECLLLEPGLNEIMANSLDYNERLWAWESWRSEVGKQLRPLY
ACE2_fish       SEMSTIYNTATVCKIDDPTDCQTLEPGLESIMAESRDYDERLHVWEGWRVATGMKMRPLY
                . *****.*..**: *:* :*  *****:.***:* **:*** .**.**  .* ::****

ACE2_HUMAN      EEYVVLKNEMARANHYEDYGDYWRGDYEVNGVDGYDYSRGQLIEDVEHTFEEIKPLYEHL
ACE2_fish       EKYVDLKNEAAKLNNYEDHGDYWRGDYETIDDPKYSYSRDQVIEDARRIYKEILPLYKEL
                *:** **** *: *:***:*********. .   *.***.*:***..: ::** ***:.*

ACE2_HUMAN      HAYVRAKLMNAYPSYISPIGCLPAHLLGDMWGRFWTNLYSLTVPFGQKPNIDVTDAMVDQ
ACE2_fish       HAYVRAKLQDVYPGHIGSDACLPAHLLGDMWGRFWTNLYPLMIPYPDRPDIDVSSAMVEQ
                ******** :.**.:*.. .*******************.* :*: ::*:***:.***:*

ACE2_HUMAN      AWDAQRIFKEAEKFFVSVGLPNMTQGFWENSMLTDPGNVQKAVCHPTAWDLG-KGDFRIL
ACE2_fish       GWDEIRLFKEAEKFFMSVNMPAMFDNFWNNSMFIKP-EERDVVCHPTAWDMGNRKDFRIK
                .**  *:********:**.:* * :.**:***: .* : :..********:* : ****

ACE2_HUMAN      MCTKVTMDDFLTAHHEMGHIQYDMAYAAQPFLLRNGANEGFHEAVGEIMSLSAATPKHLK
ACE2_fish       MCTKVNMDDFLTVHHEMGHNQYQMAYRNHPYLLRDGANEGFHEAVGEIMSLSAATPSHLQ
                *****.******.****** **:***  :*:***:*********************.**:

ACE2_HUMAN      SIGLLSPDFQEDNETEINFLLKQALTIVGTLPFTYMLEKWRWMVFKGEIPKDQWMKKWWE
ACE2_fish       SLGLLPSDFKQDYETDINFLLKQALTIVGTLPFTYMLEEWRWQVFKAKIPKDEWMQQWWQ
                *:***..**::* **:**********************:*** ***.:****:**::**:

ACE2_HUMAN      MKREIVGVVEPVPHDETYCDPASLFHVSNDYSFIRYYTRTLYQFQFQEALCQAAKHEGPL
ACE2_fish       MKRELVGVAEAVPRDETYCDPPALFHVSGDYSFIRYFTRTIYQFQFQEALCKAAGHTGPL
                ****:***.*.**:*******.:*****.*******:***:**********:** * ***

ACE2_HUMAN      HKCDISNSTEAGQKLFNMLRLGKSEPWTLALENVVGAKNMNVRPLLNYFEPLFTWLKDQN
ACE2_fish       YKCDITNSTKAGDKLRHMLELGRSMSWTRALEEVAGTTKMDSQPLLHYFSTLMEWLKEEN
                :****:***:**:** :**.**:* .** ***:*.*:.:*: :***:**..*: ***::*

ACE2_HUMAN      K--NSFVGWSTDWS-PYADQSIKVRISLKSALGDKAYEWNDNEMYLFRSSVAYAMRQYFL
ACE2_fish       QKNNRVPGWNVNVNPEISENAFKVRISLKSALGNEAYTWNANDIYLFKSTMAFAMRQYYL
                :  * . **..: .   :::::***********::** ** *::***:*::*:*****:*

ACE2_HUMAN      KVKNQMILFGEEDVRVANLKPRISFNFFVTAPKNVSDIIPRTEVEKAIRMSRSRINDAFR
ACE2_fish       KEKNTDVNFTPENIHTYNETARISFKFAVMDPTKTGTVIPKAEVENAIWQERDRINGAFL
                * **  : *  *:::. * ..****:* *  *.:.. :**::***:**  .*.***.**

ACE2_HUMAN      LNDNSLEFLGIQPTLGPPNQPPVSIWLIVFGVVMGVIVVGIVILIFTGIRDRKKKNKARS
ACE2_fish       LSDETLEFVGLMATLAPPKEEKITIWLVVFGVVMGVTVLAGIYLVTTGILNRKKKAKKAK
                *.*::***:*: .**.**::  ::***:******** *:. : *: *** :**** *  .

ACE2_HUMAN      GENPYASIDISKGENNPGFQNTDDVQTSF
ACE2_fish       EASVENPYDGSDDGEVNKAFEEDIEQTGL
                  .   . * *.. :     : *  **.:
```

**Supplemental Figure 2: Multiple sequence alignment between human ACE2 and *Danio rerio* ACE2 isoform X1.** Human ACE2 and *D. rerio* ACE2 share 58.2% primary sequence similarity. The residues highlighted in interact with the RBD domain of SARS-CoV-2 Spike protein (PDBID: 6M0J) ref: doi 10.1038/s41586-020-2180-5. *D. rerio* ACE2 has 12 similar residues out of 13 residues of hACE2 that belongs to the protein-protein interface, in which 7 of them are identical. Human ACE2 (accession code Q9BYF1), and *D. rerio* ACE2 (accession code XP_005169417). The alignment was performed using ClustalW.
